# A Signaling Hub in the Mosquito Rectum Coordinates Reproductive Investment After Blood Feeding

**DOI:** 10.1101/2025.08.25.672111

**Authors:** Chloe Greppi, Kyle Frank, Victoria Saltz, Laura B. Duvall

## Abstract

After a blood meal, female *Aedes aegypti* mosquitoes suppress host-seeking while converting ingested nutrients into yolk protein for egg development. Neuropeptide Y-like Receptor 7 (NPYLR7) is required for this behavioral switch, but its physiological role and sites of action were unknown. We identify a specialized, non-neuronal population of *npylr7*-expressing cells in the rectal pads of the mosquito hindgut. While this tissue is associated with fluid and ion balance, *npylr7* mutants maintain fluid regulation but show impaired oocyte provisioning. These cells display neuroendocrine features, including calcium responses to the NPYLR7 ligand RYamide and amino acids, as well as expression of neurotransmitter synthesis and vesicle release machinery. Vesicle recruitment occurs in these cells post-blood meal in wild-type, but not *npylr7* mutants. Our findings reveal an unexpected role for NPYLR7 in a rectal cell population senses nutritional cues and communicates with the nervous system to regulate reproductive physiology, paralleling gut-brain circuits in mammals.

## INTRODUCTION

Female *Aedes aegypti* mosquitoes require blood protein to lay eggs and successfully reproduce. Females of this species are strongly driven to bite and blood feed from human hosts and can transmit pathogens during blood feeding that cause diseases including yellow fever, Zika, dengue, and chikungunya^1–3^. The drive to blood feed is not constitutive and, after a female blood feeds to repletion, host-seeking is strongly suppressed until her eggs are laid several days later^4–10^. Behavioral shifts in responsiveness from host to oviposition-related chemosensory cues occur as females suppress their attraction to humans and begin seeking standing water for egg-laying^11^. Finding ways to exploit endogenous feeding suppression to disrupt mosquito reproduction represents an efficient approach to reduce the mosquito/human interactions that transmit disease-causing pathogens.

Neuropeptides are important regulators of feeding and are evolutionarily-conserved across phylogeny^12–15^. Decades of work have implicated Neuropeptide Y (NPY)-related signaling in the regulation of host-seeking behavior in *Aedes aegypti*. While the key C-terminal residue of mammalian NPY is tyrosine (Y), insects show diversification of NPY-related ligands, including NPF, sNPF, HP-I, and RYamide peptides^16^. Despite this diversification, the receptors that detect these ligands share approximately 60% sequence similarity with mammalian NPY receptors, and some mammalian-targeted peptides and small molecules can activate insect receptors^10,17^. Injecting peptides that activate NPY-like receptors induces host-seeking suppression in non-fed females, and genetic or pharmacological manipulation of NPY-like Receptor 7 (NPYLR7) alters host-seeking independently of blood feeding^4,10,18–20^. Although NPYLR7”s behavioral effects are established, its physiological functions and tissue(s) of action remain unclear^10^. NPYLR7 responds to multiple ligands *in vitro*, including sNPF3 and RYamide, both of which suppress host-seeking *in vivo*^10,20^. Coordination of host-seeking behavior and reproduction requires crosstalk between the nervous system, midgut, fat body, and ovaries, and relevant neuropeptides are expressed in both the nervous system and gut^21–23^. Although neuropeptide signaling that regulates behavior is typically attributed to the brain, one key NPYLR7 ligand, RYamide, is also produced by neurons that innervate the rectum^18,24^. Recent findings suggest that RYamide is released locally in the hindgut following a protein-rich blood meal, suggesting that neuropeptide signaling relevant to host-seeking suppression may occur outside of the brain^18^.

In this study we demonstrate that *npylr7* coordinates nutrient utilization and reproductive output through specialized cells in the mosquito rectum that communicate with the nervous system. Although *npylr7* mutant females consume and digest blood normally, they fail to adequately provision developing oocytes with protein. Using transcript localization and genetic reporters, we find that *npylr7* is expressed exclusively in a small population of non-neuronal epithelial cells in the rectal pads in close proximity to projections from rectum-innervating neurons originating in the ventral nerve cord. Using a genetically encoded calcium reporter, we demonstrate that *npylr7*-expressing cells respond to the application of RYamide peptide and amino acids with increases in calcium. Through RNA sequencing and immunofluorescence, we show that the hindgut expresses genes and proteins linked to neurotransmitter synthesis and vesicle release, and we identify candidate output signals from *npylr7*-expressing cells. Electron microscopy reveals that blood feeding induces recruitment of vesicle pools in these cells in wild-type females, but not in *npylr7* mutants. We propose that RYamide is locally released in the rectum from hindgut-innervating neurons in the ventral nerve cord after blood feeding, activating NPYLR7 and triggering vesicle-mediated signaling from these rectal cells back to the nervous system or peripheral tissues. This signaling axis regulates oocyte provisioning and modulates host-seeking behavior, likely through downstream effects on central circuits. These findings redefine the role of the mosquito rectum in blood feeding nutrient utilization and fecundity, uncover a previously unrecognized site of neuropeptide action critical for reproductive success, and describe an accessible target for disrupting mosquito attraction to humans.

## RESULTS

### *npylr7* is exclusively expressed in the rectal pads of the hindgut

Chemosensory-driven food seeking is regulated by NPY-related signaling in the central nervous system in many animals^25–29^. Because host-seeking and its suppression rely heavily on chemosensory modulation^6,30–34^ we initially expected that, while *npylr7* might be broadly expressed, its effects on host-seeking would likely be mediated directly by the nervous system. To characterize the anatomical expression of *npylr7* we performed Reverse Transcription-Polymerase Chain Reaction (rt-PCR) on the thorax, head and legs, and the abdomen of female mosquitoes. Unexpectedly, we detected *npylr7* transcript exclusively in samples that contained the hindgut, which is subdivided into the ileum and rectum (**Figure 1A and 1F**). Localization of transcripts specifically to this region was confirmed with fluorescence RNA *in situ* hybridization (**Figure 1B**). To identify the specific cells that express *npylr7*, we employed a genetic approach using the QF2-QUAS binary expression system to generate a “knock-in/knock-out” allele^35–37^. Insertion of the QF2 transcriptional activator under the control of the endogenous *npylr7* gene locus allows for expression of QF2 in genetically defined cells and can be combined with effectors to drive expression of a fluorescent reporter to label them (**Figure 1C**). We used CRISPR-Cas9 genome editing to make a double-stranded cut in the single exon of *npylr7* and provided a cassette for homology-directed repair containing the T2A ribosomal skipping element, the QF2 transcriptional activator, and a fluorescent reporter driven by the *Aedes aegypti* polyubiquitin enhancer/promoter to mark the insertion (**Figure 1D**). We confirmed that the transgene was integrated into the *npylr7* locus in frame and results in complete loss of *npylr7* expression when homozygosed (**Figure 1E**). To label *npylr7-*expressing cells, we crossed the *npylr7*^*QF2*^ driver line to an effector line (QUAS-CD8:GFP) to drive the expression of membrane-bound GFP in cells that express *npylr7*. In line with our rt-PCR data, we did not observe expression of *npylr7* transcript or QF2-driven reporters in the nervous system including the brain and ventral nerve cord, peripheral sensory tissues, midgut, or reproductive tissues (see **Figure S1B**). Instead, we detected CD8 and GFP signal exclusively in the rectal pads, also known as rectal papillae. The rectal pads are multi-cellular, cone-shaped structures that protrude into the gut lumen at the apical tip and make contact with the hemolymph at the basal side^38^ **(Figure 1F)**. These structures are present in many insects and are critical for fluid homeostasis and ion reuptake at both larval and adult stages, similar to the mammalian proximal colon^39–42^. A canal originating from the basal side of the structure contacts the hemolymph and houses neuronal and tracheal projections^38^. *npylr7* expression is restricted to the hemolymph-facing basal ring of each rectal pad (**Figure 1F**) and this expression is not detected in either parental line (**Figure S1A**). *Aedes aegypti* adult females have six pads, while males only have four, and we detected QF2-driven CD8:GFP signal in all four rectal pads of adult male mosquitoes (**Figure S1C**)^38^. These data indicate that *npylr7* expression is entirely restricted to the rectal pads in adult mosquitoes.

**Figure 1:**
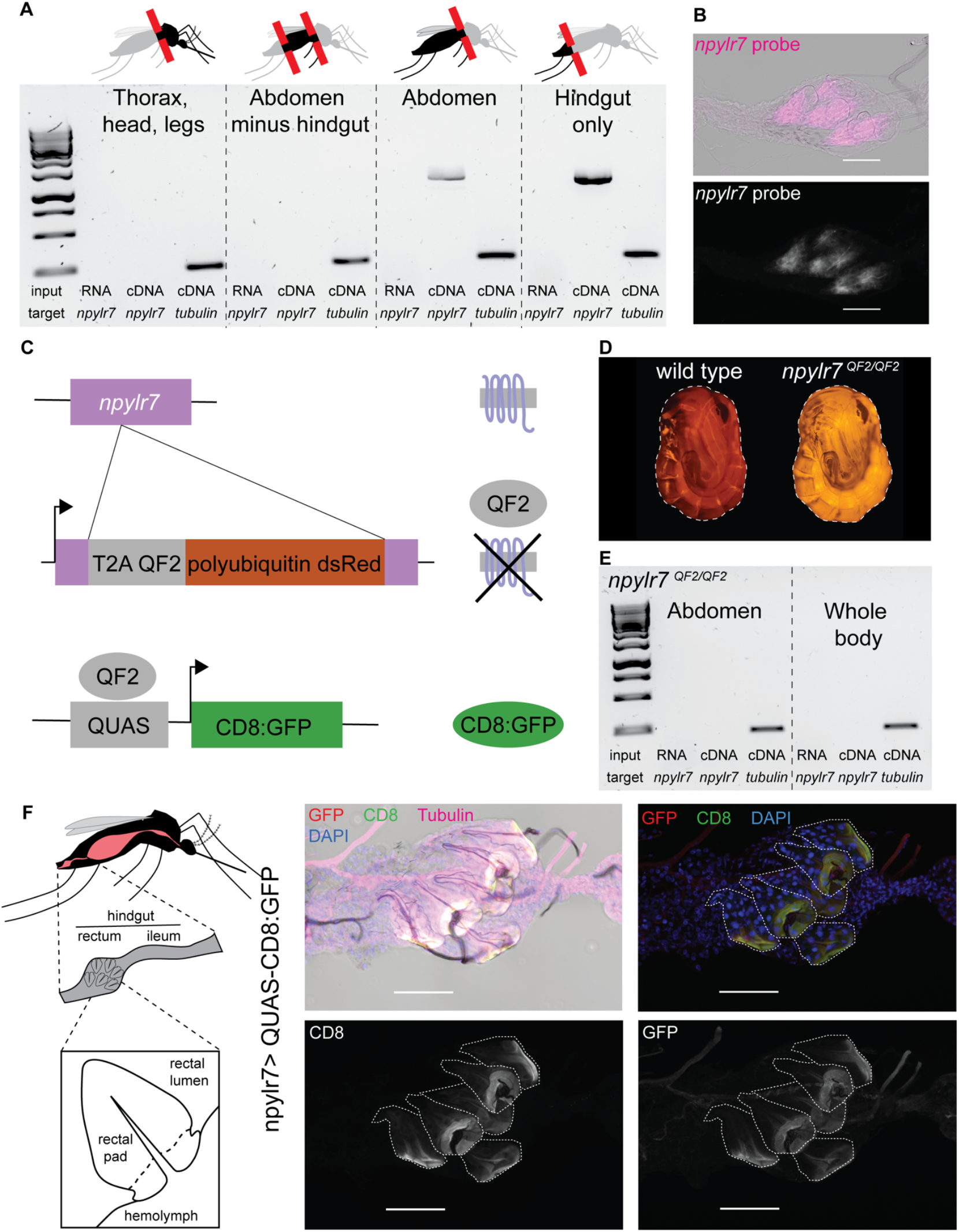
Localization and genetic manipulation of *npylr7* in *Aedes aegypti*. (A) Transcript localization using rt-PCR from mosquito tissues with primers directed against *npylr7* and *tubulin*. (B) Fluorescence *in situ* hybridization in the rectum of adult females using probes directed against *npylr7*. Scale bar = 100 µM. (C) Schematic of transgenic tools used to disrupt the *npylr7* locus and drive expression of reporters in *npylr7*-expressing cells. (D) Representative images of wild-type and homozygous mutant pupae expressing pUb-dsRed, used to mark the insertion of the transgene. (E) rt-PCR confirming loss of *npylr7* RNA transcript in homozygous *npylr7*^*QF2*/*QF2*^ animals. (F) Cartoon of rectal pads in adult mosquitoes with dashed line indicating the basal ring. Immunofluorescence of *npylr7*-QF2-driven CD8:GFP using antibodies against CD8 and GFP. Final image was stitched together from individual panels. Scale bar = 100 µM.

### Loss of *npylr7* causes deficits in egg viability and protein provisioning

Disruption of *npylr7* results in inappropriate host-seeking behavior after a blood meal during the period of oocyte maturation^10^. Because the primary function of blood feeding is to support egg development, we investigated whether this behavioral defect reflects an underlying disruption of the reproductive pathway by assessing blood-feeding efficiency and fecundity. We found that *npylr7* mutants developed normally, reached the same body weights as wild-type and heterozygous controls, and consumed blood meals of normal size (**Figure 2A**). While there was no difference in clutch sizes between genotypes (**Figure 2B**), *npylr7* mutant females had significantly reduced and highly variable egg viability (**Figure 2C**). These findings suggest that, although the mutants consumed sufficient blood nutrients to develop a fully viable clutch of eggs, they failed to effectively provision their developing oocytes to support normal hatch rates. These phenotypes were also observed in the original *npylr7* line used in a previous publication^10^, and in an independently-generated line, in which the QF2-containing cassette was inserted out of frame resulting in a disruption of the gene at a different location (**Figure S2A-C**).

**Figure 2:**
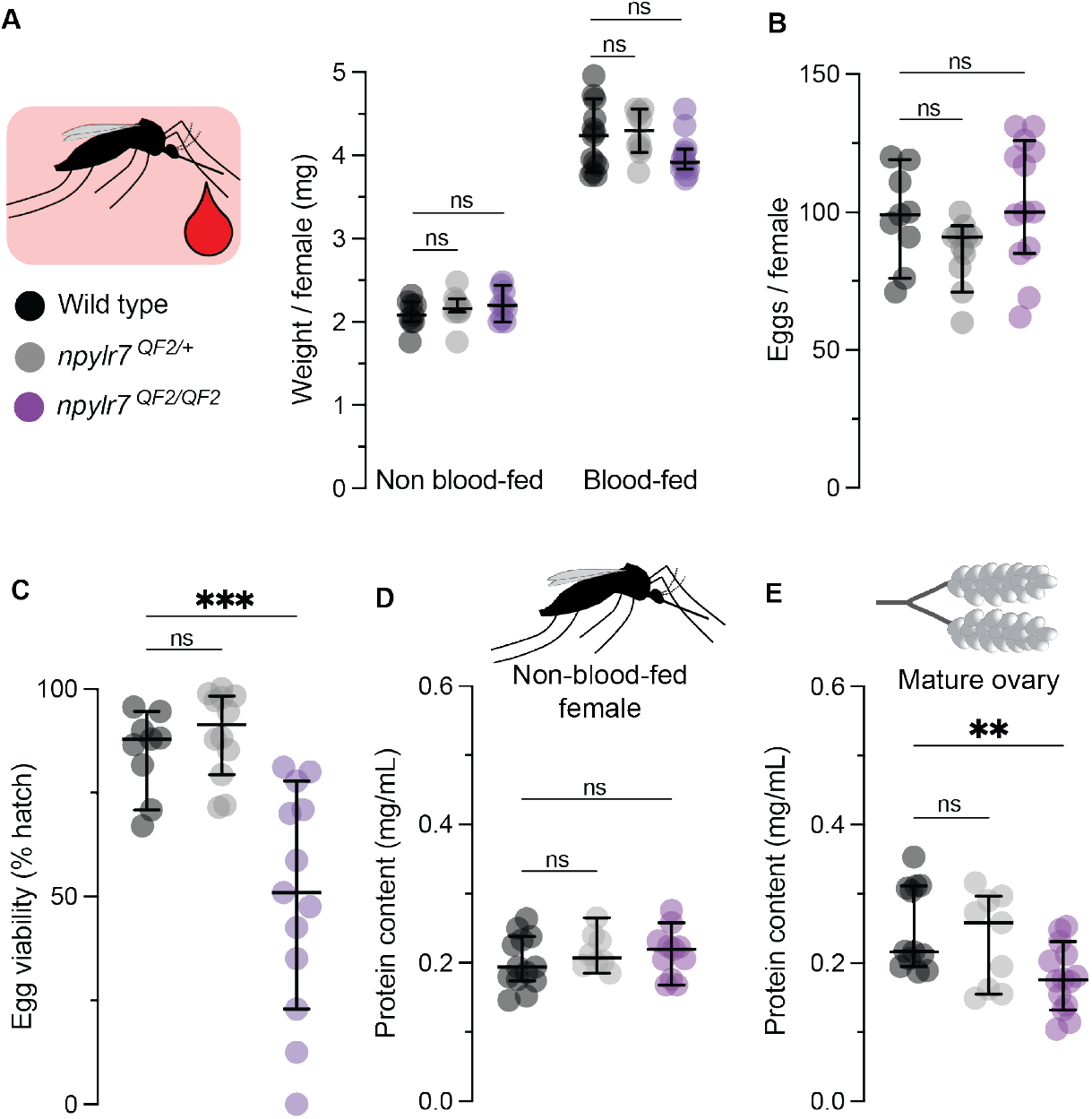
Loss of *npylr7* leads to deficits in oocyte protein provisioning and hatching. (A) Weight per female before and after blood feeding. n = 9-13 replicates (5 females per replicate) (Kruskal-Wallis test with Dunn”s multiple comparisons, ns = p > 0.05). (B) Total eggs laid. n = 9-13 clutches from individual females (One way ANOVA with Dunnett”s multiple comparisons, ns = p > 0.05). (C) Egg viability (percent of eggs hatched per female). n = 9-13 clutches (from females of 2B) (One way ANOVA with Tukey”s multiple comparisons, *** = p < 0.001). (D) Total protein content of whole females before a blood meal. n = 8-13 groups of 5 females (Ordinary one-way ANOVA with Dunnett”s multiple comparisons test, ns = p > 0.05). (E) Total protein content of ovaries 3 days post blood meal. n = 9-14 replicates of 2-3 pairs of ovaries (Kruskal-Wallis test with Dunnett”s”s multiple comparisons, ** = p < 0.01). Data are shown as medians with 95% confidence intervals for visual consistency. Statistical comparisons were performed using parametric tests for normally distributed data and non-parametric tests for non-normal data, as determined by normality test (Shapiro-Wilk). Post-hoc multiple comparison tests were conducted against the wild-type control.

After a blood meal, oocytes grow as yolk proteins are deposited over the course of 2 to 3 days^43–45^. To assess oocyte development we dissected and stained oocytes with DAPI to measure the oocyte length before and after a blood meal.

While sugar-fed, pre-vitellogenic oocytes of all three genotypes were the same size, we found that oocytes were smaller in both *npylr7* mutant lines 1 and 2 days post blood meal, during the period when most yolk protein is deposited^46^ (**Figure S2D**), suggesting that egg provisioning is abnormal when *npylr7* is disrupted. While increase in oocyte length is a commonly used proxy for yolk uptake, length is not a direct measure of protein deposition. To determine whether protein levels differ between genotypes we performed a Bradford assay to quantify total protein content of individual females before a blood meal and of ovaries 3 days after a blood meal, when yolk deposition is complete and oocytes are fully developed. Consistent with a lack of provisioning, ovaries from *npylr7* mutants had lower total protein 3 days after a blood meal, although females from all groups had similar protein content before a blood meal (**Figure 2D-E**). These data indicate that loss of *npylr7* results in deficits in utilizing blood nutrients to successfully allocate developing oocytes with sufficient protein for hatching.

### Diluting blood nutrients does not mimic loss of *npylr7*

Reduced fecundity in *npylr7* mutants could result from inefficient digestion and uptake of blood protein or a lack of available nutrients. We measured trypsin levels in the gut after a blood meal to determine whether loss of *npylr7* impaired the production of digestive enzymes. While trypsin levels were slightly higher in mutants compared to wild-type animals at 4 hours post blood meal, levels were otherwise similar, peaked 20-24 hours after a meal, and returned to pre-meal levels by 36 hours in both genotypes (**Figure S2E**). This suggests that mutant females can detect that they have consumed a blood meal and produce enzymes required to digest it appropriately. We next wondered whether additional nutrients could rescue egg viability deficits by offering *npylr7* mutant females a second blood meal 1-3 days following a first blood meal and assessing egg production and viability in females that re-blood-fed before oviposition. Consistent with the inappropriate host-seeking previously reported in *npylr7* mutants^10^, these females reliably consumed a second meal before laying their first clutch of eggs (**Figure S3A**), but this did not affect clutch size or improve egg viability (**Figure S3B-C**).

We next asked whether we could mimic the *npylr7* mutant phenotype by feeding wild-type females diluted blood meals with insufficient protein to support a full clutch of eggs. To test this, we fed wild-type and mutant females blood meals diluted 1:5 in saline. Meal size was unaffected by dilution (**Figure S3D**); however, both genotypes produced fewer eggs per female when fed diluted blood. Wild-type females laid an average of 81.67 ± 12.87 eggs per female after undiluted blood meals compared to 28.05 ± 9.15 eggs after diluted meals. Similarly, *npylr7* mutants laid 73.42 ± 16.90 eggs on undiluted blood and 34.17 ± 8.82 on diluted blood (**Figure S3E**). Wild-type females fed diluted blood also showed a modest reduction in hatch rate, 77.60 ± 12.85% compared to 88.76 ± 5.74% from females fed undiluted blood. However, this did not recapitulate the more pronounced reduction and variability in hatch rates observed in *npylr7* mutants, which were unaffected by blood meal dilution (50.59 ± 22.94% for diluted vs. 59.94 ± 25.33% for undiluted) (**Figure S3F**). These results suggest that *npylr7* mutants can detect protein content and adjust clutch size, but fail to properly provision their eggs to ensure viability regardless of meal protein content.

Thus, the reproductive defects in *npylr7* mutants are not due to impaired protein sensing or digestion, but likely reflect a failure in downstream processes that regulate oocyte provisioning.

### *npylr7* mutants show intact blood, saline, and sucrose clearance

The insect hindgut plays a key role in maintaining fluid and ion homeostasis, using cuticular pores in the rectal pads to reabsorb water, ions, and small metabolites such as amino acids, while larger waste products are excreted^39,47–49^. In *Drosophila melanogaster*, anatomical disruption of the rectal pads leads to salt-handling defects and lethality when animals are challenged with high-salt meals^50^. Blood is a physically demanding meal which presents a significant fluid regulation challenge due to its high salt content, toxic breakdown products such as heme, and large volume. To test whether *npylr7* mutants have general fluid-handling impairments, we developed assays to track the consumption and clearance of blood, saline, and sucrose meals. Multiple assays were necessary because blood and sugar feeding serve distinct physiological functions, rely on different anatomical pathways, are consumed at different volumes, and are cleared over different time courses.

Previous work suggested that *npylr7* mutants retain a portion of their blood meal longer than wild-type mosquitoes^10^. To examine whether blood excretion is affected, we developed an assay in which blood-fed females were individually housed in 6-well plates lined with filter paper. Blood excretion, visible as red spots, was quantified by measuring the area of excreta as a percentage of the total area of the filter paper. Consistent with earlier findings, *npylr7* mutants consumed normal-sized blood meals (**Figure 3A**). Both *npylr7* mutants and heterozygous animals showed more excretion events than wild-type animals on day 3, consistent with the previous report that it takes slightly longer for a blood meal to clear in *npylr7* mutants. However, blood excretion was complete by day 4 across all genotypes (**Figure 3B**).

**Figure 3:**
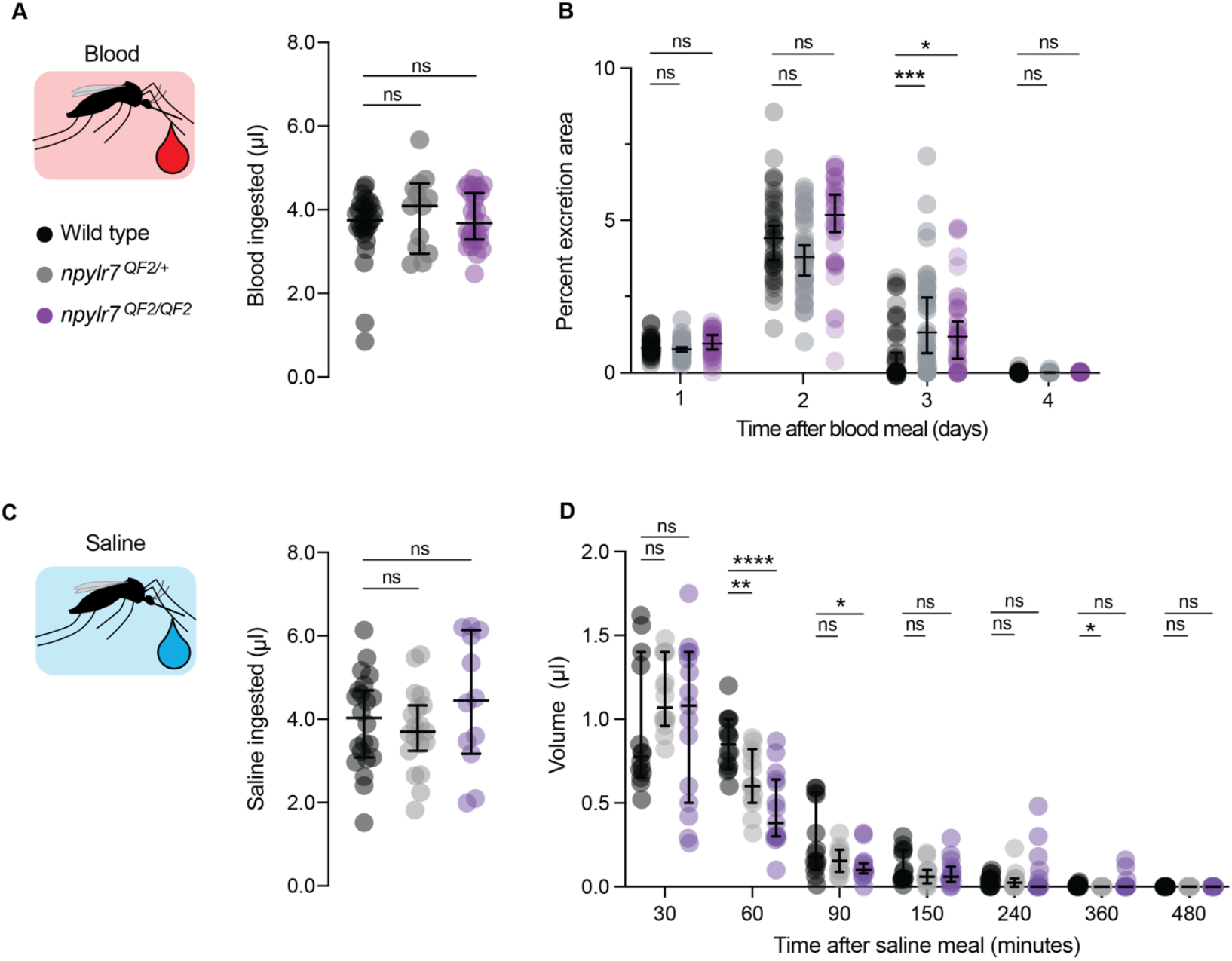
Blood and saline excretion are intact in *npylr7* mutants. (A) Blood meal size per female, n = 13-28 females (Kruskal-Wallis test with Dunn”s multiple comparisons, ns = p > 0.05). (B) Blood excretion area per female, n = 32-44 females (Kruskal-Wallis test with Dunn”s multiple comparisons, * = p < 0.05, *** = p < 0.001). (C) Saline meal size per female, n = 12-22 females (Ordinary one-way ANOVA, ns = p > 0.05). (D) Saline clearance, n = 14-15 groups of 5 females (2-way ANOVA with Dunn”s multiple comparisons test, * = p < 0.05, ** = p < 0.01, **** = p < 0.0001). Data are shown as medians with 95% confidence intervals for visual consistency. Statistical comparisons were performed using parametric tests for normally distributed data and non-parametric tests for non-normal data, as determined by normality test (Shapiro-Wilk). Post-hoc multiple comparison tests were conducted against wild-type control.

To test fluid regulation independently of digestion, we fed females a saline meal isotonic with blood and supplemented with ATP. Although this non-nutritive meal does not support egg development, it engages the “blood feeding” program and induces engorgement^51,52^. After feeding, we housed groups of 5 females in packed cell volume tubes and quantified excretion volume at different time intervals. All groups consumed saline meals of comparable volume (**Figure 3C**), and, although mutants showed reduced clearance at the 60 and 90-minute timepoints (**Figure 3D**), all genotypes cleared this meal in the expected time frame of 6-8 hours^53^ and showed no lethality in the days following saline ingestion.

Mosquitoes also consume sugar meals, which are much smaller in volume than blood meals and used primarily for energy homeostasis, as opposed to egg development^43^. To determine whether *npylr7* mutants showed general deficits in consumption or excretion of a sugar meal, females were provided access to a fluorescein-labeled meal of 10% sucrose and collected at intervals after feeding. Sugar intake and clearance were then quantified by fluorescence measurements. Although we observed some variation between wild-type and heterozygous animals, mutants showed no deficits in sucrose meal size (**Figure S4A**) and were able to clear their sugar meal within 48 hours (**Figure S4B**), confirming that sugar handling remains intact.

Although we observed some variation in clearance rates between genotypes, all meals were ultimately cleared within the expected timeframe in all genotypes without lethality. This suggests that the core functions of the rectal pads in fluid and ion homeostasis remain intact in *npylr7* mutants and raises the possibility that this tissue serves roles beyond fluid regulation.

### *npylr7*-expressing cells are non-neuronal, but the rectum is highly innervated

Our data indicate that *npylr7* function in the rectal pads is associated with protein provisioning after a blood meal as opposed to controlling fluid handling. To investigate how NPYLR7 may be activated after a nutritive blood meal has been consumed we anatomically characterized the innervation patterns of the mosquito rectum using an established pan-neuronal QF2 line to express CD8:GFP^54^. Although the rectal pads themselves are non-neuronal, they are densely innervated with extensive processes originating from neurons with cell bodies in the ventral nerve cord (**Figure 4A**). These projections extend through the basal ring into the central canal and closely appose *npylr7*-expressing cells, suggesting the potential for direct communication (**Figure 4B**). Recent work has shown that neuronal projections to the central canal express RYamide and suggests that this neuropeptide is released from these projections within hours of the ingestion of protein-rich meals^18,24^. Given that RYamide is a known activator of NPYLR7 *in vitro*^10^, it is a strong candidate for mediating direct activation of NPYLR7 in the rectum.

**Figure 4:**
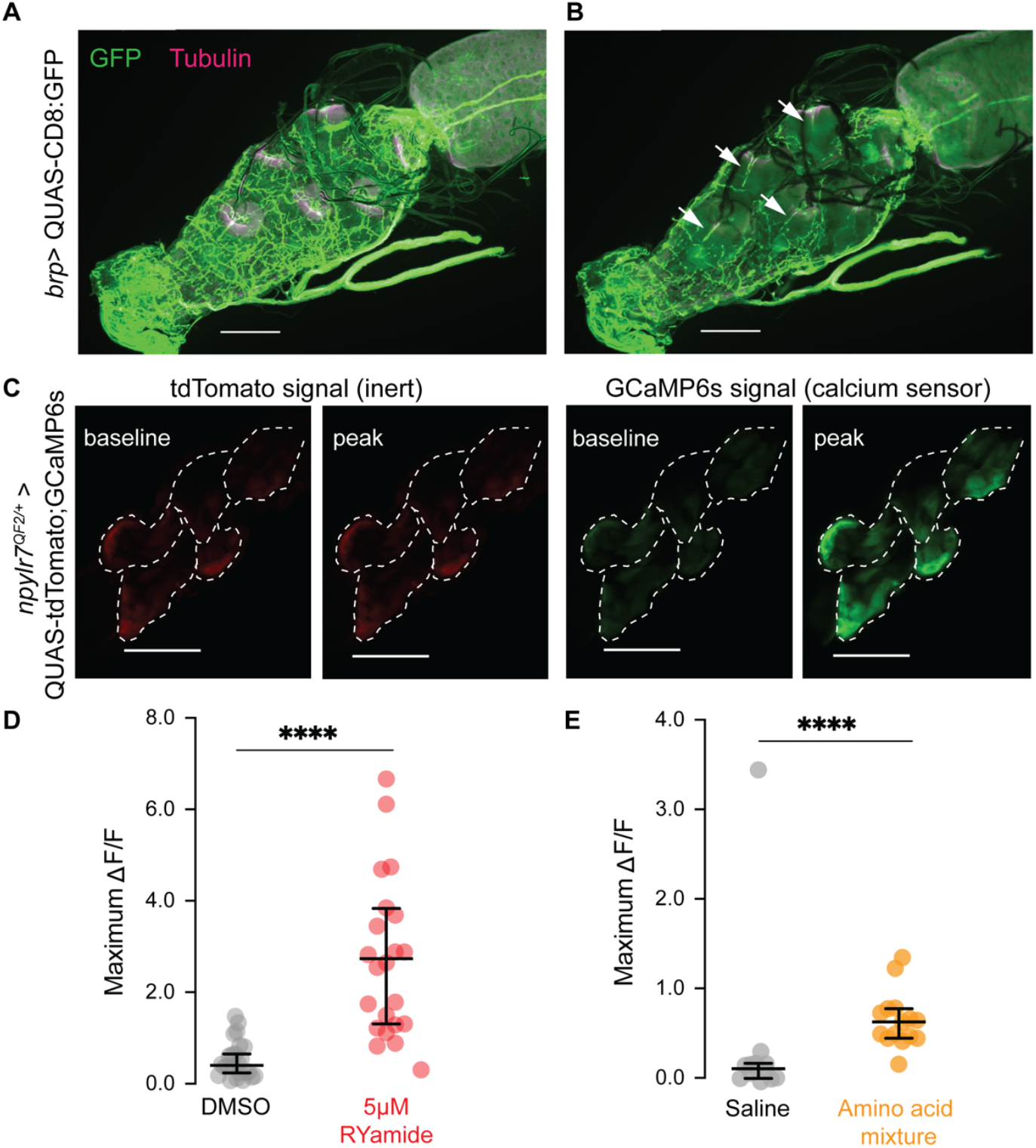
*npylr7* cells are highly innervated and respond to RYamide and amino acid application with calcium influx. (A) *Bruchpilot (Brp)* positive innervation of the surface of the rectum from neurons originating in the ventral nerve cord. Final images were stitched together from individual panels. Scale bars = 100 µm. (B) Neuronal projections that enter the central channels of the rectal pads and closely appose *npylr7*-expressing cells, indicated with white arrowheads. Final images were stitched together from individual panels. Scale bars = 100 µm. (C) Images of tdTomato and GCaMP signal at baseline and maximal response to application of 5µM RYamide peptide. Dashed lines indicate individual rectal pad boundaries. Scale bars = 100 µm. (D) Maximum change in fluorescence over baseline of DMSO controls and RYamide, n = 22-24 ROI from 10-14 females (Mann-Whitney, **** = p < 0.0001). (E) Maximum change in fluorescence over baseline of DMSO controls and amino acid mixture, n = 13-15 ROI from 10-11 females (Mann-Whitney, **** = p < 0.0001). Data are shown as medians with 95% confidence intervals for visual consistency. Statistical comparisons were performed using parametric tests for normally distributed data and non-parametric tests for non-normal data, as determined by normality test (Shapiro-Wilk).

### *npylr7*-expressing cells respond to RYamide and amino acid application

To determine whether *npylr7*-expressing cells can directly respond to RYamide stimulation, we used a functional imaging approach. By driving the calcium sensor GCaMP6s in *npylr7*-expressing cells using our *npylr7*^*QF2*^ heterozygotes, we monitored calcium activity in an *ex vivo* rectal pad preparation. Bath application of 5 μM RYamide triggered a rapid and significant increase in GCaMP fluorescence compared to DMSO controls, demonstrating that *npylr7*-expressing cells in the rectal pads respond to RYamide with calcium increases (**Figure 4C–D** and **Figure S5A**).

Although these cells are hemolymph-facing, their location in the rectum may also expose them to signals from the gut lumen. Previous RNA sequencing analysis detected expression of amino acid transporters and sensors in the hindgut^55^ (see also **Data S1**), prompting us to ask whether *npylr7*-expressing cells also respond to amino acids. We found that application of an amino acid mixture, at concentrations that mimic human blood, induced increases in GCaMP fluorescence, while amino acid-free saline controls did not (**Figure 4E** and **Figure S5B**). These responses were weaker in amplitude compared to those elicited by RYamide and may occur through indirect activation, with similarities to glutamine responses reported in the mouse colon^56^. Although our *ex vivo* preparation does not allow us to distinguish between luminal and hemolymph-derived sources of amino acids, our findings demonstrate that *npylr7*-expressing rectal cells can respond to both peptide and nutrient cues. This suggests they may serve as sensory effectors, integrating postprandial signals from the hemolymph and potentially the gut lumen to regulate processes involved in protein allocation following blood feeding.

### The mosquito hindgut expresses transcripts associated with neuroactive signaling

While transcriptional studies of the mosquito gut have revealed region-specific functions that vary by meal type, the rectum has often been excluded from these analyses either due to dissection protocols or its traditionally understood role in resorption^55^.

However, our findings suggest that the rectum also engages neuroactive signaling, including pathways downstream of neuropeptide receptor activation. To identify candidate genes that could mediate this signaling we performed RNA sequencing on 80-100 pooled hindguts from non-blood-fed female mosquitoes. This unbiased transcriptomic approach provides a comprehensive view of gene expression in the hindgut, allowing us to identify both established and novel components of its gene expression repertoire. To preserve tissue integrity, dissections included both the ileum and rectum. All RNA samples had high quality (mean quality score greater than 38) and 77–81% of reads mapped uniquely to the genome (AaegL5.0 assembly). As expected, we observed expression of genes corresponding to Gene Ontology (GO) term categories that were uniquely associated with the hindgut in previous studies^55^, including transcripts associated with fluid homeostasis, ion reuptake, and amino acid transport (**Figure 5A**). However, we also identified GO terms related to synaptic transmission, neurotransmitter release, and neuropeptide signaling (**Figure 5A**). Specifically, we detected transcripts associated with secretion, neurotransmitter biosynthesis, and synaptic machinery. In addition to *npylr7* (LOC5570400), we found expression of other neuropeptide receptors known to regulate diuresis and other processes in the rectal pads, including Kinin receptor (LOC5574266) and Pyrokinin Receptor 1(LOC5576812)^49,57^ (**Figure 5D**).

**Figure 5:**
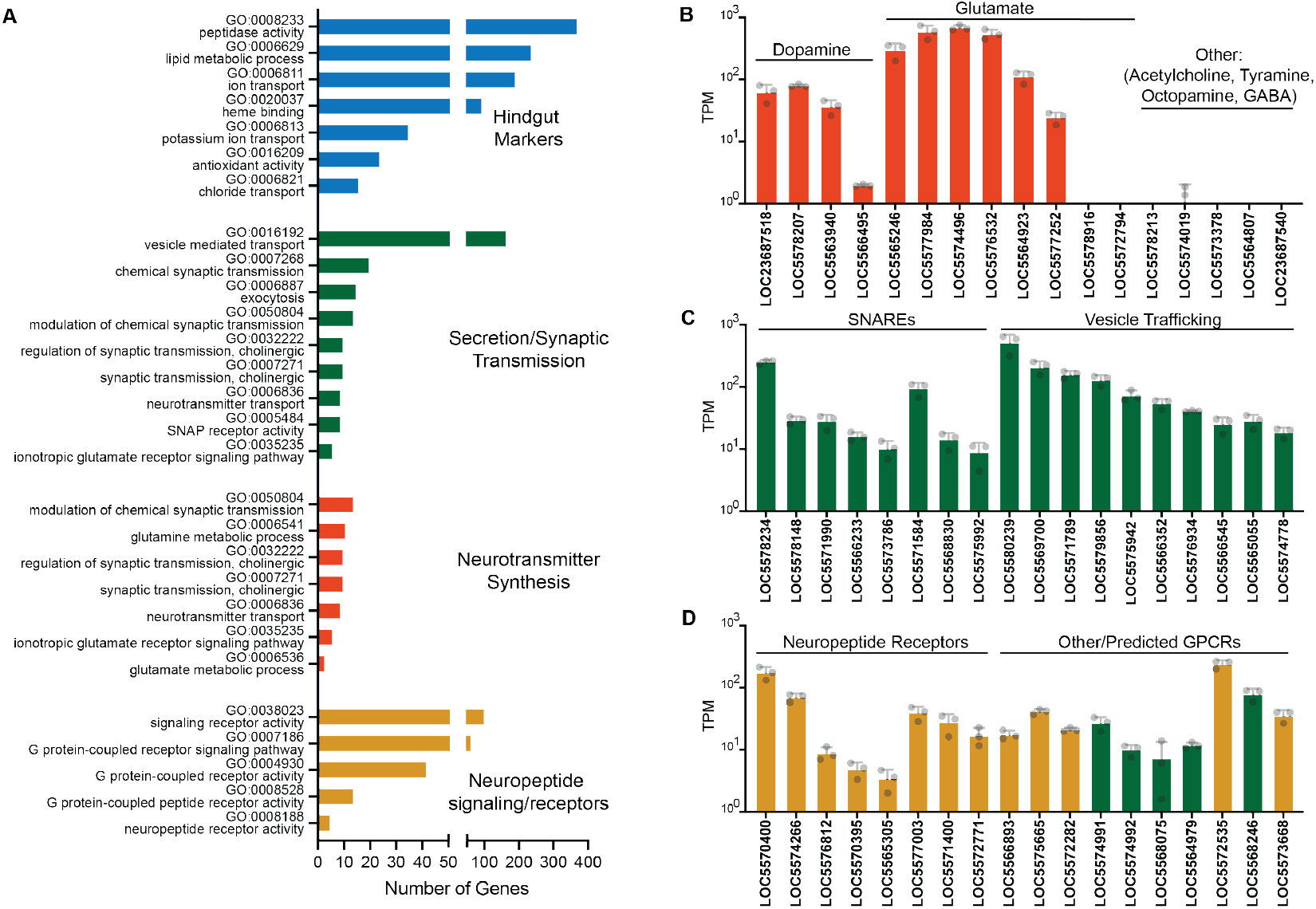
Expression of transcripts in the mosquito ileum and rectum. (A) GO terms associated with hindgut markers^55^ and secretion, synaptic transmission, and neuropeptide signaling and receptors. (B) Expression levels of enzymes involved in neurotransmitter synthesis in the mosquito hindgut. (C) Expression levels of SNAREs and vesicle release components in the mosquito hindgut. (D) Expression levels of neuropeptide and neurotransmitter receptors in the mosquito hindgut.

Strikingly, we found significant expression of enzymes required for synthesizing glutamate and dopamine but not for GABA, acetylcholine, octopamine, or serotonin (**Figure 5B**). Although the rectum is densely innervated by projections from neurons originating in the ventral nerve cord we did not observe neuronal cell bodies in this region (see **Figure 4A**), nor did we detect appreciable levels of the canonical neuronal marker *brp1*^58^, making it unlikely that the observed transcripts arise from nearby neurons (**Data S1**). While GO analysis flagged cholinergic signaling terms, closer inspection revealed that the annotated genes were either uncharacterized or not part of canonical cholinergic pathways. We also found transcripts encoding SNARE proteins and other components of vesicular release machinery (**Figure 5C**), suggesting that this tissue may be capable of regulated secretion.

Together, these findings indicate that the mosquito hindgut may play an active signaling role through the synthesis and release of neuroactive compounds including dopamine and glutamate, an unappreciated function for this tissue.

### *npylr7* cells co-express secretory machinery and enzymes for glutamate synthesis

The dense innervation of the rectal pads may provide inputs to *npylr7*-expressing cells through the release of RYamide and other signals, but they may also function to receive signals released locally from these cells and relay them back to the nervous system. The hemolymph-facing position of *npylr7*-expressing cells places them in an ideal location to secrete short-range signals to adjacent neuronal projections or to release long-range signals that may act hormonally. To test whether these cells possess the machinery for neurotransmitter synthesis and release, we examined co-expression of secretory components and glutamate synthesis enzymes detected in our bulk sequencing using immunofluorescence and fluorescent *in situ* hybridization (FISH). Immunostaining revealed that glutamine synthetase is co-expressed with *npylr7*-driven CD8:GFP, as well as with Synaptotagmin 1, a marker of secretory vesicles expressed throughout the pads (**Figure 6A**). FISH further confirmed the presence of syntaxin and glutamate synthase transcripts in the rectal pads, including the basal ring where *npylr7* is expressed (**Figure 6B-C**). These data support the idea that *npylr7*-expressing cells are equipped to produce and secrete neurotransmitters.

**Figure 6:**
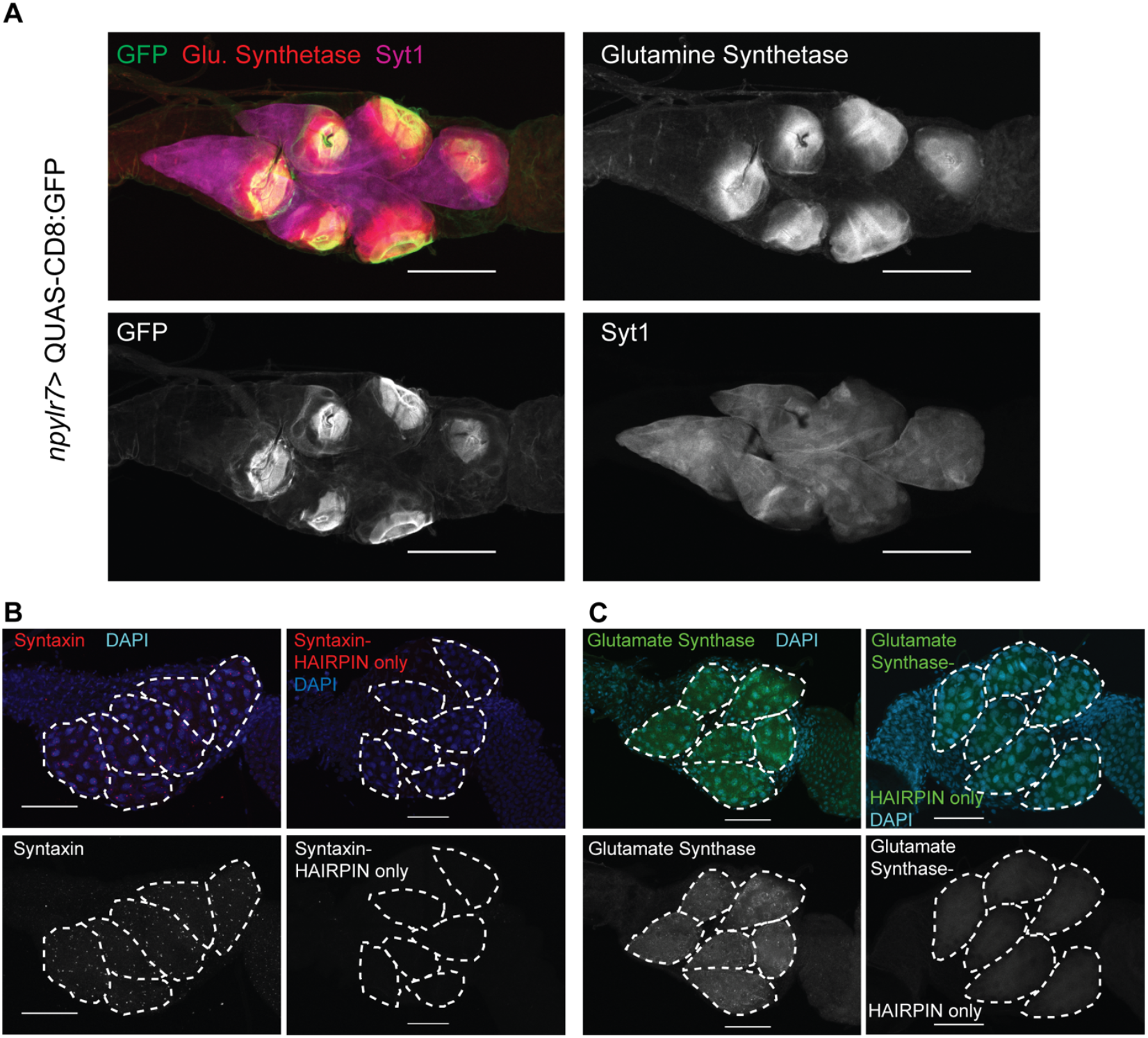
Rectal pads express transcripts and proteins associated with neurotransmitter synthesis and vesicle release. (A) Immunostaining with antibodies against synaptotagmin (magenta) and glutamine synthetase (red) with *npylr7*-driven CD8:GFP (green). (B) RNA-FISH with syntaxin (red), DAPI (blue) and hairpin only control. (C) RNA-FISH with glutamate synthase (green), DAPI (blue) and hairpin only control. Final images were stitched together from individual panels. All scale bars = 100 µm.

### Vesicle pools accumulate on the basal side of the rectal pads following a blood meal

Although the presence of transcripts and proteins associated with synaptic function suggests the potential for neuroactive signal release, these markers do not provide direct evidence of vesicle secretion. To investigate whether *npylr7*-expressing cells exhibit ultrastructural features associated with vesicle release, we used electron microscopy to visualize the rectal pads before and 6 hours after a blood meal. This timepoint aligns with the proposed release of RYamide peptide and putative NPYLR7 activation^18^. Consistent with prior studies, in non-blood-fed wild-type females we observed an organized, mitochondria-rich ultrastructure^38^ with isolated vesicular structures approximately 35 nm in diameter (**Figure 7A**). Within 6 hours of a blood meal, organized pools of vesicles were recruited specifically to the basal side of the rectal pads, the region where *npylr7* is expressed (**Figure 7B**). We observed clear structural differences in *npylr7* mutants compared to wild-type females even in the non-blood-fed state with a lack of visible organization of vesicular structures and the presence of multilamellar structures that were rarely observed in wild-type females (**Figure 7C**). These structures are similar to multilamellar bodies that have been previously identified in invertebrate digestive tissues^59,60^ and may be associated with heavy metal stress^61^. Most strikingly, mutant females completely failed to recruit vesicle pools after blood feeding, and we observed an increase in the presence of multilamellar bodies (**Figure 7D**). These data provide evidence that *npylr7*-expressing cells structurally remodel after blood feeding. The recruitment of vesicle pools suggests that these cells are poised to release chemical signals after a blood meal and this remodeling is dependent on *npylr7*.

**Figure 7:**
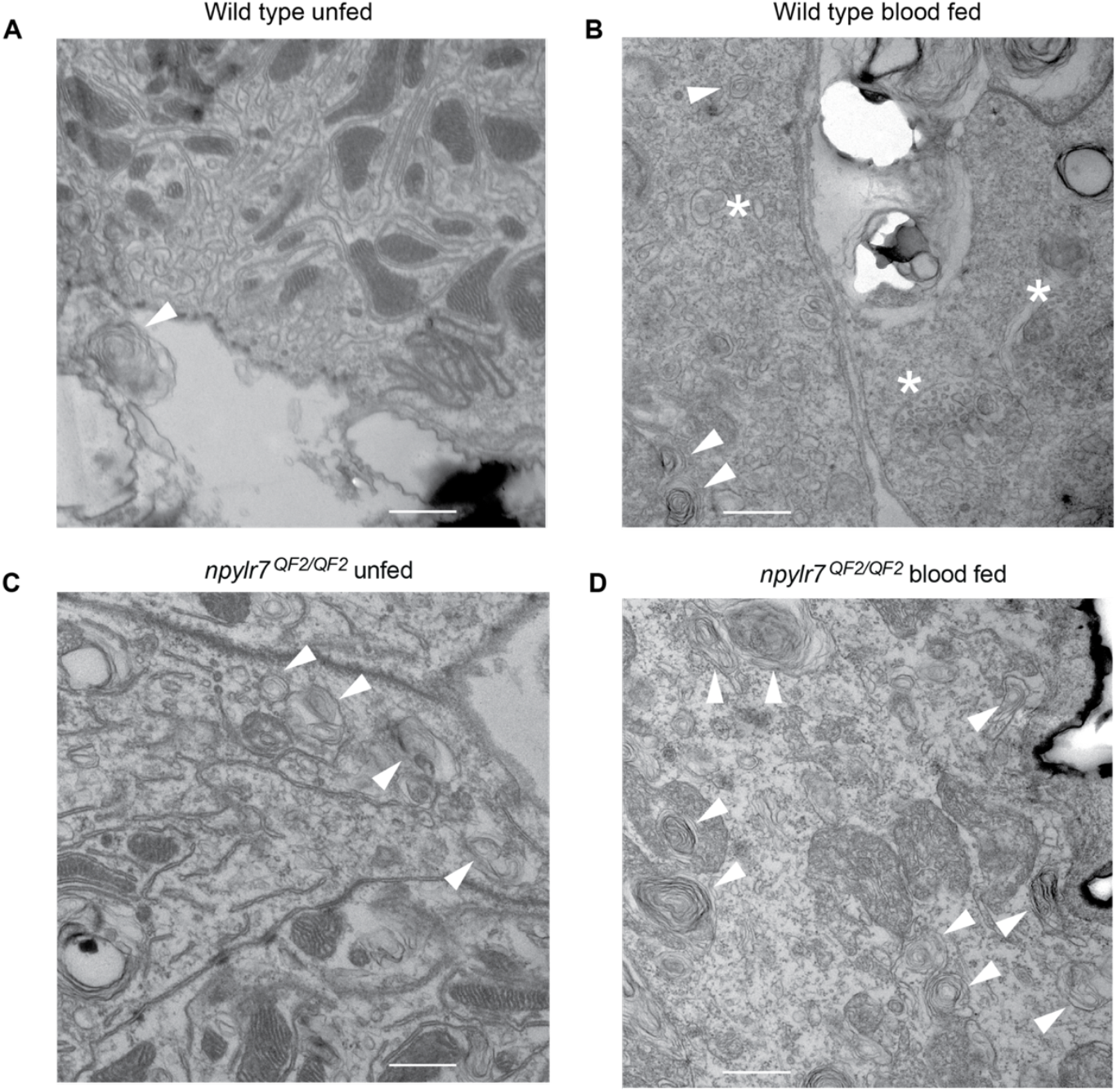
Vesicle pools are recruited to the basal side of the rectal pads 6 hours after a blood meal in wild-type, but not *npylr7* mutant females. (A) EM of wild-type non-fed female rectal pads (Imaged at 23,000x). (B) EM of wild-type female rectal pads, 6-hours post blood meal (Imaged at 30,000x). (C) EM of *npylr7*^*QF2/QF2*^ non-fed female rectal pads (Imaged at 30,000x). (D) EM of *npylr7*^*QF2/QF2*^ female rectal pads, 6-hours post blood meal (Imaged at 30,000x). All scale bars = 500 nm, arrowheads highlight multi-lamellar structures and asterisks label vesicle pools. **Supplemental Information**

## DISCUSSION

Our work uncovers a novel role for a rectum-expressed neuropeptide receptor, NPYLR7, in coordinating reproductive physiology in *Aedes aegypti*. While NPYLR7 has been established as a key regulator of host-seeking behavior, we show that it is critical for appropriate oocyte protein provisioning. This receptor is found in a highly restricted set of cells on the basal side of the rectal pads in the hindgut that respond to peptides and amino acids and express markers of neuroactive signaling. Located at the interface between the digestive tract, hemolymph, and nervous system, these *npylr7* cells are well positioned to integrate physiological signals and modulate reproductive output and behavior.

### Linking hindgut signaling to oocyte provisioning

Although the insect hindgut is primarily associated with fluid and ion balance, our findings reveal its unexpected involvement in nutrient allocation during oogenesis. Despite its localization in the hindgut, *npylr7* is dispensable for the clearance of blood, saline, and sucrose meals. Instead, mutant females exhibit impaired oocyte protein provisioning, despite normal feeding behavior and early digestive processes. Protein levels in mature oocytes and hatch rates are reduced, though not abolished, suggesting that *npylr7* is not strictly essential but serves a modulatory “quality control” role, possibly to set protein thresholds for allocation. While most nutrient transport occurs in the midgut, our transcriptomic data and recent studies reporting amino acid transporter expression in the hindgut^55^ suggest that this tissue may participate in nutrient uptake or sensing. One hypothesis is that RYamide activation of NPYLR7 following a blood meal triggers protein uptake from the hindgut, serving as a secondary or “last chance” mechanism to ensure that sufficient resources reach the developing oocytes. Alternatively, rectal pad sensing of amino acid levels could influence nutrient absorption elsewhere, for example via feedback to the midgut. The premature return to host-seeking observed in *npylr7* mutants^10^ may reflect a physiological response to under-provisioned oocytes, which raises the intriguing possibility that reproductive tissues signal unmet nutrient demands to drive feeding behavior.

### Mechanisms of sensing and signaling from the hindgut

Among previously-identified NPYLR7 ligands, RYamide is a strong candidate for activating this receptor *in vivo*. This neuropeptide is produced by abdominal neurons that innervate the rectal pads and is likely released within hours of blood feeding^18,24^. Although the rectal pads project into the gut lumen, *npylr7*-expressing cells are positioned basally facing the hemolymph, an efficient location to sense systemic cues. Consistent with activation by local peptide release, *npylr7*-expressing cells show calcium increases upon RYamide application.

Although these cells do not express classical neuronal markers including *bruchpilot*, our RNA-seq analysis confirmed the presence of G proteins in the hindgut (**Data S1**), but whether NPYLR7 activation triggers calcium influx directly via G protein signaling or indirectly through other extracellular or intracellular sources remains unclear. Beyond peptide responses, *npylr7* cells also exhibit modest calcium responses to amino acids, suggesting a nutrient-sensing function that may be mediated independently of NPYLR7 activation. Whether these nutrient cues originate from the gut lumen or hemolymph remains unresolved using our *ex vivo* preparations, but these cells are well-positioned to integrate both sources.

Morphological and transcriptional evidence supports a secretory role for *npylr7*-expressing cells. Electron microscopy showed that after a blood meal, these cells accumulate pools of vesicles averaging 35 nm diameter, consistent with small clear vesicles that typically store classical neurotransmitters.

Notably, these vesicle pools are absent in blood-fed *npylr7* mutants, supporting the importance of NPYLR7 signaling in post-feeding vesicle recruitment. RNA-seq and immunofluorescence suggest these cells may be capable of synthesizing and releasing dopamine and glutamate. We detected transcripts for VMAT, which is needed for dopamine packaging, but not vGLUT, which typically loads glutamate into vesicles. However, glutamate could be released via vGLUT-independent mechanisms, such as reverse transport by excitatory amino acid transporters or ATP-gated purinergic channels^62,63^.

Because our profiling was performed on non-blood-fed females it is possible that additional neurotransmitters are synthesized and released after feeding. We also detected transcripts for key enzymes for neuropeptide processing (**Data S1**), suggesting that neuropeptides may be part of their secretory output. Understanding how these cells are activated by peptide and nutrient cues, identifying the anatomical sources of these signals, and mapping their signaling outputs and downstream targets will be critical to understand how these rectal cells coordinate inter-organ communication and drive postprandial physiological and behavioral responses.

### Comparisons with other insects

The behavioral sequence of host seeking, blood feeding, and digestion requires coordinated communication across multiple organs, yet until now, the hindgut has not been implicated in this process. Our findings expand the scope of gut-brain communication in insects and highlight the diverse roles of neuropeptide signaling beyond classical sites. While neuropeptides can modulate sensory circuits via direct effects on neurons in the brain^25^, our results suggest that NPYLR7 signaling in *Aedes aegypti* operates through a rectum-brain axis.

Interestingly, RYamide-expressing neurons also innervate the rectal pads in *Drosophila melanogaster*, though RYamide appears to be evolving into a pseuodogene in that species and produces little to no peptide^64^. This raises the possibility that RYamide signaling has been co-opted or expanded in blood-feeding mosquitoes to regulate reproduction-linked feeding behavior. Investigating whether other hematophagous insects express RYamide in abdominal neurons and cognate receptors in the rectal pads may reveal conserved or convergent evolution of this signaling axis.

### Parallels to mammalian gut–brain signaling

In mammals, NPY receptors and peptide ligands are broadly distributed across the central nervous system and gastrointestinal tract and play critical roles in feeding regulation through their actions in central and peripheral tissues^65–69^. In contrast, *Aedes aegypti npylr7* exhibits an unusually restricted expression pattern, limited to a small population of rectal cells. The NPY Y2 receptor, which is most similar to *Aedes aegypti* NPYLR7, is enriched in the colon and involved in regulating motility, satiety, and electrolyte absorption^68,70–76^. Peptide YY (PYY), a gut-derived ligand of Y2R, is released postprandially and modulates feeding behavior, gut motility, and nutrient processing^77,78^. PYY has also been shown to increase water and electrolyte absorption in the ileum^79,80^ reinforcing the idea that gut-localized neuropeptide signaling can modulate both physiological and behavioral responses to feeding.

There is precedent for non-neuronal, electrically excitable cells in the mammalian gut that communicate directly with the nervous system. Many features of *npylr7*-expressing cells resemble specialized neuroendocrine cells described in the mammalian gut including mammalian L cell, enterochromaffin, or “neuropod” cells that can form synapse-like contacts with neurons and release neuroactive compounds^81–84^. These specialized cells express synaptic markers such as synapsin-1, release glutamate, and respond to glucose with calcium transients^81,82^. Their synaptic connections with the vagus nerve enable rapid signaling from the gut to the brain about nutrient identity and availability. Similarly, *npylr7*-expressing cells may be capable of releasing neurotransmitters that can rapidly relay information back to the ventral nerve cord and ultimately to the brain to link postprandial nutrient status and the regulation of reproductive feeding behaviors.

### Implications for vector control

Understanding how feeding behavior and reproductive physiology are linked offers opportunities for novel mosquito control strategies. NPYLR7 represents an attractive target because its activation suppresses host-seeking, while its loss impairs reproductive output. In addition, the expression of this receptor in the gut makes it accessible to pharmacological modulation via ingestion^10,85^. Compounds that activate NPYLR7 could prolong post-feeding satiety, reducing biting frequency and pathogen transmission. Conversely, antagonists could disrupt reproductive success.

More broadly, targeting gut-based signaling hubs that integrate feeding, nutrient sensing, and reproduction may provide powerful new tools for mosquito control and disease prevention.

## RESOURCE AVAILABILITY

Further information and requests for reagents should be directed to and will be fulfilled by the lead contact, Laura B. Duvall. All data and code are available in **S1 data table**.

## Supporting information

DataS1

## ACKNOWLEDGEMENTS

We thank Thomas Gabel for assistance with animal husbandry, Isabelle Seckler for assistance with genotyping and homozygosing mutants, Margo Herre for assistance with the HCR protocol, and Andrew Matheson for guidance on RNAseq analysis. Electron microscopy was performed at the University of Colorado, Boulder EM services Core Facility in the MCDB Department, with technical assistance by Garry Morgan and Sarah Zimmermann. We thank Nilay Yapici, Helen Cox, Oliver Hobert, Leslie Vosshall, and members of the Duvall lab for comments and useful discussions on the manuscript. This work was supported by the following grants: NIGMS (R35 GM137888) (LBD), Beckman Young Investigator Award (LBD), Pew Scholar in Biomedical Sciences Award (LBD), Klingenstein-Simons Fellowship Award in Neuroscience (LBD), LSRF Postdoctoral Fellowship (CG).

## AUTHOR CONTRIBUTIONS

**C.G**.: Conceptualization, Methodology, Validation, Formal analysis, Investigation, Resources, Writing-Original Draft, Writing-Review & Editing, Project Administration, Funding acquisition. **K.F**.: Methodology, Validation, Investigation, Writing-Review & Editing. **V.S**.: Methodology, Validation, Investigation, Writing-Review & Editing. **L.B.D**.: Conceptualization, Methodology, Formal analysis, Resources, Writing-Original Draft, Writing-Review & Editing, Supervision, Project Administration, Funding acquisition.

## SUPPLEMENTAL INFORMATION

Supplemental Figure and Raw Data files, GEO (Gene Expression Omnibus NIH) https://www.ncbi.nlm.nih.gov/geo/info/submission.html

## EXPERIMENTAL MODEL DETAILS

### Mosquito rearing and maintenance

The following strains (*Aedes aegypti* Liverpool background) were used in this study: wild-type *Aedes aegypti* Liverpool, *npylr7*^*QF2*^ (this study), *npylr7*^*3xP3*^ (this study), QUAS-mCD8-GFP^86^, brp-QF2 and QUAS-GCaMP6s^54^. All mosquito lines were maintained and reared at 28°C, 70-80% relative humidity with a 12 h light: 12 h dark schedule.

Mosquito eggs were hatched using 1 L of deoxygenated water containing 1 TetraMin fish food tablet. Larvae were fed TetraMin tablets as needed. Adults were maintained in cages with *ad libitum* access to 10% sucrose solution. Female mosquitoes used for experiments were between 5 and 15 days old and co-housed with males. For passaging, mutant generation, and blood feeding assays, adult female mosquitoes were fed defibrinated sheep blood (QuadFive) with 1 mM ATP at 37°C using a Hemotek artificial membrane feeder system. For all feeding assays (blood, saline and sucrose) animals were starved overnight by removing sucrose but providing access to water to prevent desiccation.

### Generation of *npylr7*-QF2 knock-in knock-out line

The *npylr7*^*QF2*^ line was generated using established methods^37^. Three sgRNAs were designed using FlyCRISPR (https://flycrispr.org/) to target *Aedes aegypti* Neuropeptide Y-like Receptor 7 (AAEL019786) using the AaegL5 reference genome. Efficiency of the sgRNAs was evaluated by injecting sgRNA and Cas9 protein into 200 *Aedes aegypti* Liverpool embryos. The synthesis of the sgRNA and the preparation of an injection mix was performed as described^37^: DNA oligonucleotide templates were amplified using the KOD polymerase and purified with NucleoSpin columns. T7 Megascript Kit was used for *in vitro* reverse transcription. RNA was purified using MegaClear column purification kit.

Guide RNA (40 ng/µL at final concentration) was purified by ethanol precipitation and mixed with Cas9 protein (300 ng/µL at final concentration). Guide RNA with the most efficient cutting rate (TTCATCGCGAACCTCGCCCTCGG, PAM sequence underlined) was selected to design a plasmid for homology-directed repair (HDR). The transgene insert contains the left homology arm (234 bp upstream of cut site) and followed by the ribosomal skipping element T2A, transcriptional activator QF2, polyubiquitin promoter, fluorescent marker dsRed and the right homology arm (907 bp directly after the cut site). This 4,880 bp genetic insert was synthesized and cloned into a pUC57 plasmid (GenScript). The donor plasmid (700 ng/µL) was mixed with guide RNA (40 ng/µL) and purified by ethanol precipitation before mixing with Cas9 protein (300 ng/µL) and injected into ~ 1,000 *Aedes aegypti* Liverpool embryos. All embryo injections were performed by the Insect Transformation Facility at the University of Maryland Institute for Biosciences & Biotechnology Research. Injected animals were outcrossed to wild-type virgin males/females in families of < 25 injected animals to screen for germline transmission of the red polyubiquitin marker under a Leica MZ10 F fluorescence stereomicroscope.

One independent line (*npylr7*^*3xp3*^) that disrupted the *npylr7* locus but did not yield QF2 expression was isolated and maintained under standard mosquito rearing conditions. The mutation was characterized by Sanger sequencing which showed integration of the genetic insert into the designated cut site and confirmed gene disruption. All mutant lines were backcrossed to wild-type Liverpool mates for five generations before being used for experimentation. All heterozygous mutants used in this study were generated by continuing backcrossing to wild-type Liverpool animals. Offspring were screened as larvae or pupae by the presence of the red fluorescent marker under a Leica MZ10 F fluorescence stereomicroscope. To generate and enrich for homozygous mutants, heterozygous *npylr7*^*QF2/+*^ mutants were co-housed to produce a transgene-enriched population. Homozygous animals from this population were genotyped by performing PCR amplifications on DNA extracted from single legs of each virgin adult mosquito using Phire Tissue Direct PCR Master Mix following the manufacturer protocols. A primer pair (F: TTCACTGCCGAGTTCTTCCC; R: TACTGGCATCACTCTTCCGC) flanking the insertion was used, and the presence of the long, insertion-containing product and the absence of the short wild-type product was used to identify homozygous individuals. Homozygous individuals were pooled to establish homozygous lines. These lines were confirmed by the presence of the ds-Red polyubiquitin marker in all offspring when crossed to wild-type Liverpool animals.

## METHOD DETAILS

### rt-PCR

Tissues were collected from 7-20 day old animals of relevant genotype. 5-6 animals were used for abdomen or thorax/head samples and 45-50 animals for hindgut only samples. All were dissected in PBST under RNase-free conditions and on separate dissection plates. RNA was isolated using TRIzol/chloroform extraction, followed by TURBO DNase treatment for 30 minutes at 37°C, and a second TRIzol/chloroform extraction. Precipitated RNA was dried and resuspended in 20 µL of RNase-free water. RNA concentration and purity (260/280 ratio) were measured using a NanoDrop spectrophotometer. All samples were diluted to 40 ng/µL using RNase-free water. cDNA synthesis was performed on 80 ng (2 µL) of purified RNA extract using a ProtoScript II First Strand cDNA Synthesis Kit. PCR was performed using a Phire Tissue Direct PCR Master Mix with 1 µL of the cDNA product (or RNA extract) per reaction (npylr7 primers: F: TTCACTGCCGAGTTCTTCCC; R: TACTGGCATCACTCTTCCGC; tubulin primers (LOC5577489): F: CTGCTTCAAAATGCGTGAAT R: GGTTCCAGATCGACGAAA). 10 µL of PCR product from each reaction was run on 1% agarose gel and imaged using SYBR safe dye.

### Immunofluorescence

Unless otherwise noted, all immunofluorescent staining was conducted on co-housed female mosquitoes 5-15 days old. Mosquitoes were cold anesthetized on ice and hindgut was dissected in ice cold 1x PBS with 0.25% Triton-X 100 (PBST) and moved to 1 mL of fixative (4% formaldehyde in PBST) and nutated for 3 hours at 4°C. Tissues were washed five times for 10 minutes in wash buffer (PBST) and placed in blocking buffer (2% normal goat serum and 4% Triton-X in PBS) at 4°C nutating overnight. Tissue was washed three times for 10 minutes wash buffer, then placed in primary antibody solution for 1-3 days at 4°C. Primary antibody solutions were made in antibody buffer (2% normal goat serum and 0.25% Triton-X in PBS), with the following dilutions: goat anti-CD8 1:100, chicken anti-GFP 1:1,000, mouse anti-tubulin 1:100, rabbit anti-glutamine synthetase 1:250, mouse anti-Syt1 1:200. After primary antibody incubation, tissue was then washed 5 times for 20 minutes each in wash buffer before overnight incubation at 4°C in secondary antibody solution. All secondary antibody solutions were made in antibody buffer at 1:500 dilutions.

DAPI was added to secondary antibody step at 1:1,000 dilution (stock at 1 mg/mL). Tissue was washed three times for 10 minutes each in wash buffer and then mounted in Vectashield medium. Samples were kept at 4°C before imaging on a Zeiss LSM880 laser scanning confocal microscope unless otherwise noted.

### RNA-FISH

RNA in situ hybridization (HCR) were performed using reagents and protocols from Molecular Instruments (Los Angeles, CA). Custom probes were designed for the following genes: npylr7 (accession#: XM_021839074.1), syntaxin1 (accession#: XM_021840067.1), glutamate synthase (accession#: XM_021841419.1). Co-housed female wild-type mosquitoes 5-15 days old were dissected in ice-cold nuclease-free PBST and transferred to a fixative solution (4% formaldehyde and 0.25% Triton-X in PBS) and nutated at room temperature for 3 hours.

Animals were washed five times with nuclease-free wash buffer (Nuclease-free wash buffer, 0.25% Tween-20 in PBS) and transferred to 100% methanol at −20°C for 1-2 hours for dehydration. Tissue was then re-hydrated for 5 min at room temperature in 75%, 50% and 25% methanol solutions (diluted in NF wash buffer). Tissue was then washed 4 times for 5 min in 100% NF wash buffer before being transferred to 400 µL probe hybridization buffer (Molecular Instruments) and incubated at 37°C for 30 minutes. Tissue was then transferred into 250 µL of pre-heated probe hybridization buffer with 20 pmol of custom-made probe and incubated with rotation overnight at 37°C. On the following day, probes were washed with 400 µL probe wash buffer (Molecular instruments) for five times at 10 minutes each at 37°C, and then washed twice, 10 minutes each, with fresh 5x SSCT buffer (5x SSC and 0.1% Tween 20) at room temperature. Brains were then transferred into 400 µL of amplification buffer (Molecular Instruments) and incubated for 10 minutes at room temperature before incubation overnight in 100 µL amplification buffer with 9 pmol snap-cooled hairpins with rotation at room temperature. Tissue was washed once for five minutes, twice for 30 minutes, and once at five minutes again before mounting in SlowFade Diamond mounting media. Samples were kept at 4°C before imaging on a Zeiss LSM880 laser scanning confocal microscope.

### Fecundity and hatching assay

Females were blood fed defibrinated sheep blood using a Hemotek artificial feeder. Females were allowed 60 minutes to feed to repletion and females were visually scored for complete engorgement and placed in a separate cage with *ad libitum* sucrose. 3 days after the blood meal, individual females were placed in ovivials (fly vials (VWR #75813-164) with 5 mL of diH2O and a filter paper cone (Cytiva #1001-055) as oviposition substrate) and given 3 days to lay eggs. Filter paper was removed, photographed and eggs were manually counted. Egg papers were then dried for 5 days in small cups and then allowed to hatch for 3 days in de-oxygenated water. Larval counts were performed manually 4 days after hatching.

### Blood meal quantification

Blood meal quantification was performed as previously described^10^. Briefly, co-housed female mosquitoes 5-15 days old were starved overnight and fed defibrinated sheep blood supplemented with 2% fluorescein (final concentration of 0.002% fluorescein) and 1 mM ATP at 37oC using a Hemotek feeder. Prior to feeding, at least 10 unfed animals were reserved for unfed controls (3 replicates) and standard curve (7 reference points), and 100 µL of blood meal was saved for standard curve. Animals were visually screened for abdominal distension and frozen at −20°C if not processed immediately. Individuals were placed in 1.5 mL tubes with 120 µL of 1x PBS and homogenized with a pestle. A standard curve using non-fed mosquitoes and fluorescein blood meal was prepared with 0.0, 0.5, 1.0, 2.0, 3.0, 4.0, 5.0 µL reference points.

Samples were spun at 14,000 RPM for 10 min and 50 µL was loaded into each well of a 96-well plate. Fluorescence intensity (excitation: 475, emission: 510) was measured on a Biotek Synergy Neo plate reader. The standard curve was used to calculate the volume of blood ingested.

### Quantification of meal sizes by weight

Females were weighed in groups of 5 on an analytical balance (Fisher Scientific #01-913-930) before or after a blood meal fed using a Hemotek feeder. Total weights were then divided by 5 and reported as weight per female.

### Egg development quantification

Females of the relevant genotype were starved overnight and blood fed defibrinated sheep blood using a Hemotek feeder. Blood-fed females were sorted visually and kept in cages with access to *ad libitum* sucrose. 72 hours (3 days) after the blood meal, oocytes were dissected and fixed as in the immunohistochemistry protocol. After fixing, oocytes were washed 5 times for 10 min in PBST and incubated in blocking buffer with DAPI at 1:1000 dilution nutating overnight at 4°C. Oocytes were washed and mounted in Vectashield and imaged on Nikon Ti2-E microscope using 10x objective. Oocyte lengths were obtained by measuring the vertical distance in microns from the apical to basal end of the primary egg chamber using Nikon Elements software. 5-10 oocytes were measured per ovary, for a total of 10-20 oocytes per female.

### Refeeding

Females were fed sheep blood using the Hemotek feeder as described previously. Fully engorged females were collected and separated into cages designated for a second feed on either day 1, 2 or 3 after their initial meal. One control group of animals was never offered a second meal. Animals were offered a second meal of sheep blood containing 0.001% fluorescein on the designated day. Animals were cold anesthetized and scored as having taken a second blood meal by screening for fluorescent signal in the abdomen using a GFP filter under Leica MZ10 F fluorescence stereomicroscope. Animals were placed in ovivials and eggs were collected, manually counted and hatched to score fecundity as in the fecundity assay.

### Dilute blood feeding

Females were offered sheep blood diluted in saline (110 mM NaCl, 20 mM NaHCO_3_) with 1 mM ATP. Fully engorged females were aspirated and placed into new cages with access to 10% sucrose. 3 days after feeding females were placed in oviposition vials for egg laying. Eggs were then counted manually and hatched to score fecundity. Larval counts were performed manually 4 days after hatching.

### Saline excretion assay

Females were starved overnight and fed a meal of saline (110 mM NaCl and 20 mM NaHCO_3_) with 1 mM ATP and red food coloring (McCormick Red Food Color) the following day using a Hemotek feeder^87^. Animals were weighed before and after feeding and groups of 5 fed females were placed in individual packed cell volume tubes covered in thin layer of parafilm to prevent evaporation (Sigma-Aldrich #Z760986). Tubes were placed on a rack in the insectarium at 28°C, 70 – 80% relative humidity. At designated timepoints, tubes were removed from racks and briefly placed on ice to knock out animals and allow transfer to a new labeled PCV tube for the next time point. After animal removal PCV tubes were immediately spun down on a tabletop centrifuge to collect excreted saline in the bottom capillary and the volume was visually assessed and recorded. Volume excreted at each time point was divided by the number of females in the tube to record an approximate volume excreted in µL / female.

### Sugar meal quantification

Females were starved overnight (with water provided) 48 hours before the experiment, with water removed for the last 24 hours (no access to sucrose or water in the last 24 hours). For each cage, a fresh sucrose meal consisting of 25 mL of 10% sucrose, 2 drops of red food coloring (McCormick) and a final concentration of 0.002% fluorescein was prepared. Prior to feeding, dead animals were removed and 10 unfed animals were reserved for negative controls and standard curve preparation. After removal of 1mL of the sucrose meal for the standard curve, a large cotton ball was dipped into the remaining meal such that the top was saturated but the bottom remained dry to reduce drownings and placed at the bottom of the cage. Animals were given 1 hour to feed after which all mosquitoes were aspirated and placed in individual 1.5 mL tubes with 120 µL of 1x PBS. Samples were homogenized, prepared, and read on Biotek plate reader as in the blood quantification assay. The standard curve was prepared using 0.0, 0.5, 1.0, 1.5, 2.0, 2.5 and 3.0 µL known volumes of labeled sucrose meal as reference points. 3 unfed controls were included in each replicate and used for normalization.

### Sugar meal clearing

Animals were starved and fed using the same protocol as the sugar meal quantification assay. After 1 hour of feeding, the cotton ball was removed (alternatively, animals were gently aspirated and transferred to a new cage). Females were not provided water or sucrose for the remainder of the assay, while males were provided water only.

Animals were aspirated at desired time points and placed in 120 µL of 1x PBS in 1.5 mL tubes. Animals were then processed as in the blood and sugar quantification assays.

### Blood excretion assay

To measure the area of blood excretion every 24 hours after blood feeding females were individually housed on top of absorbent filter paper in 12-well tissue culture plates. 12-well plates (CELLTREAT #229111) were prepared by adding 1 mL of agarose solution (1% agarose, 10% sucrose dissolved in water) to each well and cooled overnight to ensure the females retained access to sucrose and water. Females were fed sheep blood with 1 mM ATP using the Hemotek feeder system. 12 filter paper (Cytiva #1001-125) rounds of 2.5 cm (created using a 2.5 cm stamp (Amazon B093ZXG5X7)) were placed on the lid corresponding to the well locations. Engorged females were briefly cold anesthetized, placed individually on 11 of the 12 filter papers and enclosed by placing the bottom of the plate (containing the wells) on top of the lid such that the agar-filled wells are above the mosquitoes. One well contained only the paper as a blank control. The plates were kept in the insectary throughout the assay. Every 24 hours after feeding females were briefly cold anesthetized and transferred to a freshly prepared plate with new filter papers. Filter papers were imaged using a Coomassie gel filter on gel imaging system (Bio-Rad #QQ284921-CPQ22) at a 0.046 second exposure. All images from all genotypes and timepoints were converted to an 8-bit tiff and thresholded identically to highlight excreta as black pixels. The same circular ROI (slightly smaller than the filter paper) was used to calculate the percent area of excretion. Background from the 12th blank filter paper was subtracted from all values to obtain a final percent area of excretion. All genotypes were run in parallel.

### Bradford assay

For sugar fed whole female protein levels, groups of 5 females were homogenized in 500 µl of cold 1x PBS. For ovaries 72 hours after blood feeding, females were fed defibrinated sheep blood using the Hemotek feeding system. Engorged females were reared and at 72 hours post meal their ovaries were dissected out in ice cold 1x PBS in groups of 3-5 females. Ovaries were homogenized in 500 µl of cold 1x PBS. All samples were then centrifuged at 12,000 RPM at 4oC for 20 min. Samples were processed using Bradford Assay kit (Bio-Rad, #5000204).

Briefly, 5 µl of sample was added in wells of a 96-well plate in triplicate and 250 µl of Bradford reagent was then added. The plate was shaken for 1 min and 5 min later absorbance was read at 595 nm. A standard curve provided in the kit was used to calculate the protein quantity in each sample, using the average absorbance of the 3 biological replicates. The calculated protein quantity was then divided by the number of females from that sample tube to obtain the protein concentration per female or per ovary pair.

### RNAseq sample preparation

Samples were prepared established protocols^55^. Briefly, 80-100 hindguts encompassing the ileum and rectum were dissected from females aged 5-15 days old into ice cold 1x PBS with 0.025% Triton. Samples were homogenized for 30-60 seconds in 100 µl of TRIzol reagent, then topped off with 500 µl TRIzol for a total of 600 µl. 120 µl of chloroform were added to samples, vortexed and centrifuged for 15 min at 12,000g at 4°C. 200 µl of supernatant was transferred to a fresh tube. All steps afterward used RNeasy kits (Qiagen, #74104). 700 µl of Buffer RLT and 500 µl of 100% ethanol were added, sample was vortexed briefly and run through provided columns with one final spin to fully dry the column. Columns were washed twice using 500 µl of Buffer RPE with a final spin to dry the column. Samples were eluted in 30 µl of RNase-free water. All tubes, reagents and tools were RNase free. Three replicates were prepared and sent for sequencing.

### RNAseq library preparation

RNA samples were sent to Azenta Life Sciences (South Plainfield, NJ) for library prep, sequencing, and analysis. Total RNA samples were quantified using Qubit 3.0 Fluorometer (Life Technologies, CA) and RNA integrity was checked using Agilent TapeStation 4200 (Agilent Technologies, CA).

Sequencing libraries were prepared using NEBNext Ultra II Kit for Illumina using manufacturer”s instructions. Library quality was validated using the same methods as total RNA validation. Sequencing libraries were sequenced on the Illumina NovaSeq instrument using a 2x 150bp Paired End configuration.

### RNAseq sequencing and data analysis

Sequence reads were trimmed using Trimmomatic v.0.36 and mapped to Aedes aegypti reference genome (AaegL5) available on ENSEMBL using STAR aligner v.2.5.2b to generate BAM files. Unique gene hit counts were calculated by using feature Counts from the Subread package v.1.5.2. Only unique reads that fell within exon regions were counted. After extraction of gene hit counts (including TPM) GO term analysis was performed using the g:Profiler online tool^88^ to extract GO terms of interest.

### Electron Microscopy

Hindguts (non-fed or 6 hours post blood meal of both wild-type and *npylr7*^*QF2/QF2*^ females) were dissected from 5-10 day old cold anesthetized females into cold 1x PBS. Samples were fixed overnight at 4ºC in 4% paraformaldehyde in PBS and then washed three times over 24 hours in 1x PBS and stored overnight at 4ºC prior to shipping. Samples were shipped in 1x PBS on cold packs to the BEMS facility at the University of Colorado (Boulder, CO). Samples were exchanged into 0.1 M sodium cacodylate, and then post-fixed with 1% osmium tetroxide in 0.1M sodium cacodylate for 1 hr. Post-fix media was washed twice with buffer and twice with dH2O, followed by an ethanol dehydration series over a period of 4 hrs. Ethanol was exchanged with anhydrous acetone. Resin (Epon/Araldite) infiltration occurred over 4 days at room temperature and then polymerized for 48 hours at 65ºC. Thin sections (60-80 nm) were cut using a Leica UC6 ultramicrotome, collected on Formvar-coated TEM slot grids, and poststained with 2% aqueous uranyl acetate and Reynold”s lead citrate. TEM images were collected on a Tecnai T12 Spirit TEM, operating at 100 kV, with an AMT CCD (digital camera).

### Calcium imaging

*npylr7*^*QF2*^ and QUAS-GCaMP6s animals were crossed to obtain *npylr7*^*QF2*^>QUAS-GCaMP6s females. 5-21 day old females were briefly cold anesthetized and the hindgut was dissected in ice cold saline (103 mM NaCl, 3 mM KCl, 5 mM TES, 4 mM MgCl2, 1.5 mM CaCl2, 25 mM NaHCO3, 1 mM bNaH2PO4, 10 mM Trehalose, 10 mM Glucose, pH 7.3) and placed in 1.9 mL of ice cold saline in poly-lysine coated dishes (Cellvis #d35-14-1.5-n). Rectal pads were visualized using a 20x objective, with mCherry and GFP channels adjusted for optimal adjusted gain to obtain a median brightness to allow for fluorescence fluctuation without loss and then untouched for the remainder of the assay. A 30 second baseline was obtained, after which 100 µL of 50 *µM* RYamide (for a final concentration of 5 µM) or DMSO was added. Final concentration of DMSO (0.05%) was the same for both DMSO and RYamide experiments. Responses were recorded for 3 minutes following stimulus and sampled at 1 frame per second. For amino acid responses, samples were prepared using saline without amino acids. Imaging paradigm was the same as with RYamide, including a 30 second baseline and 3 min post-stimulus recording. 100 µL of a 10x amino acid solution reflecting amino acid concentrations found in human blood and required for oocyte development was used as stimulus (final concentration: 42 *µM* Isoleucine, 100 *µM* Arginine, 87 *µM* Leucine, 185 *µM* Lysine, 65 *µM* Phenylalanine, 120 *µM* Threonine, 180 *µM* Valine, 60 *µM* Tryptophan)^89^. 100 µL of amino acid-free saline was used as the negative control.

For analysis of RYamide, amino acid responses, ROIs were drawn around the rectal pads using mCherry channel and the mean intensity measured for each frame in both GFP and mCherry channels (210 frames, 1 frame/second). GFP mean intensity was normalized to mCherry mean intensity. The average normalized GFP intensity for frames 0-30 was taken as the baseline and used to calculate the change in baseline fluorescence for frames 31-210. Max ΔF was the maximum change observed for frames 31-210. All image processing was done in FIJI and live imaging acquisition was done on a Zeiss LSM880 laser scanning confocal microscope.

## QUANTIFICATION AND STATISTICAL ANALYSIS

All statistical analyses were performed with GraphPad Prism version 10.5. Data are shown as medians with 95% confidence intervals for visual consistency unless noted otherwise. Statistical methods and sample sizes are included in figure captions. Nonparametric tests were used for data that does not follow normal distribution as determined by Shapiro-Wilk test. Data used for all figures, along with statistical test details including exact p-values, can be found in **Data S1**.

## ADDITIONAL RESOURCES

Hindgut RNA-seq data have been deposited at GEO accession#: GSE301574 and are publicly available. Original gel and microscopy data reported in this paper will be shared by the lead contact upon request. All raw data reported in the paper are available in **Data S1**.

---

**Figure S1:**
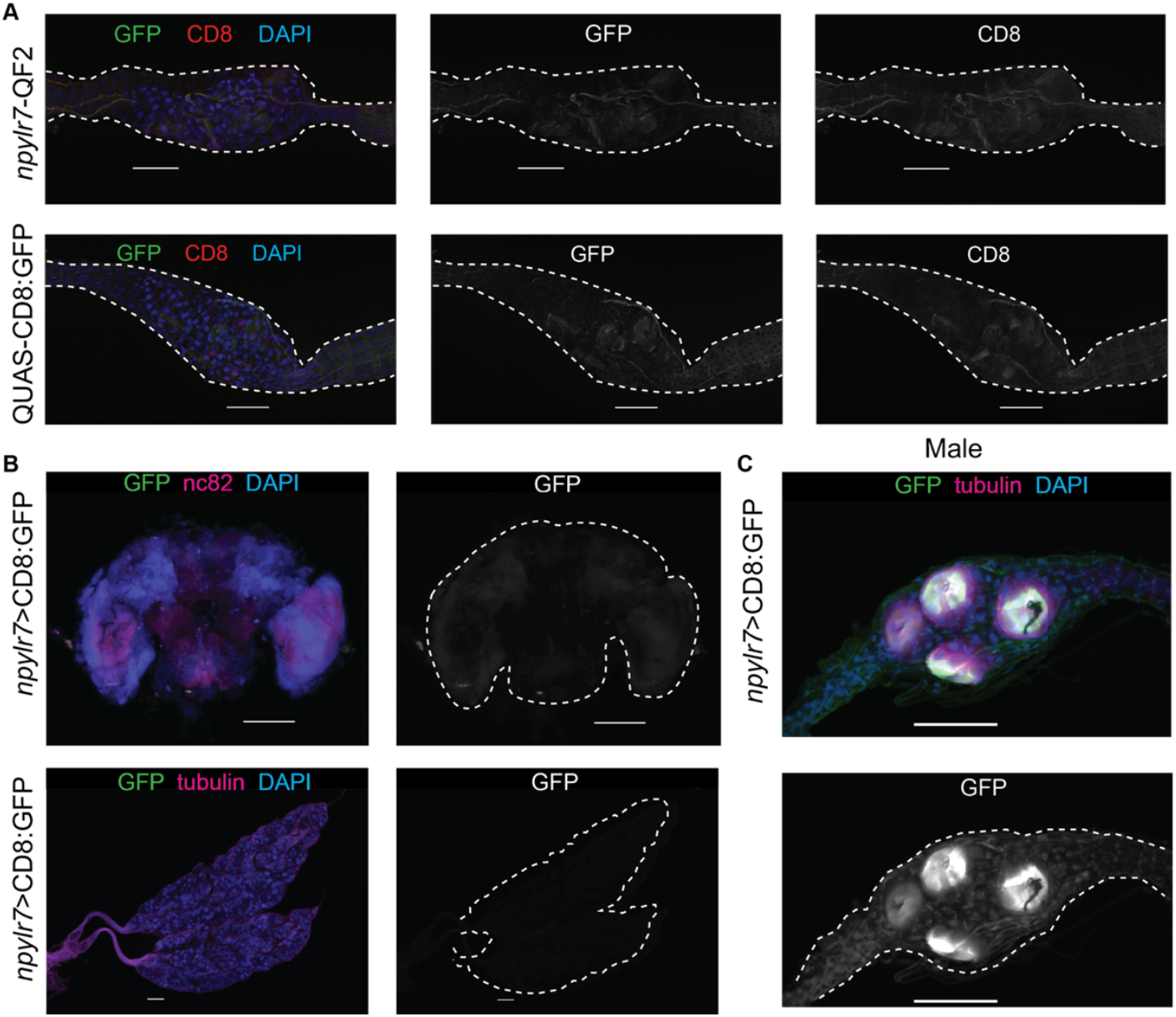
Analysis of *npylr7* expression across tissues and in male hindgut. (A) Immunohistochemical staining using antibodies against CD8 and GFP in rectum of parental controls expressing either QF2 or QUAS elements alone. (B) Immunohistochemical staining using antibodies against CD8 and GFP in the brain (top panels) or the ovary (bottom panels) of *npylr7*>CD8:GFP-expressing females. (C) Immunohistochemical staining using antibodies against CD8 and GFP in the rectal pads *of npylr7*>CD8:GFP expressing males. All final images in this figure were stitched together from smaller individual panels. All scale bars = 100 µM.

**Figure S2:**
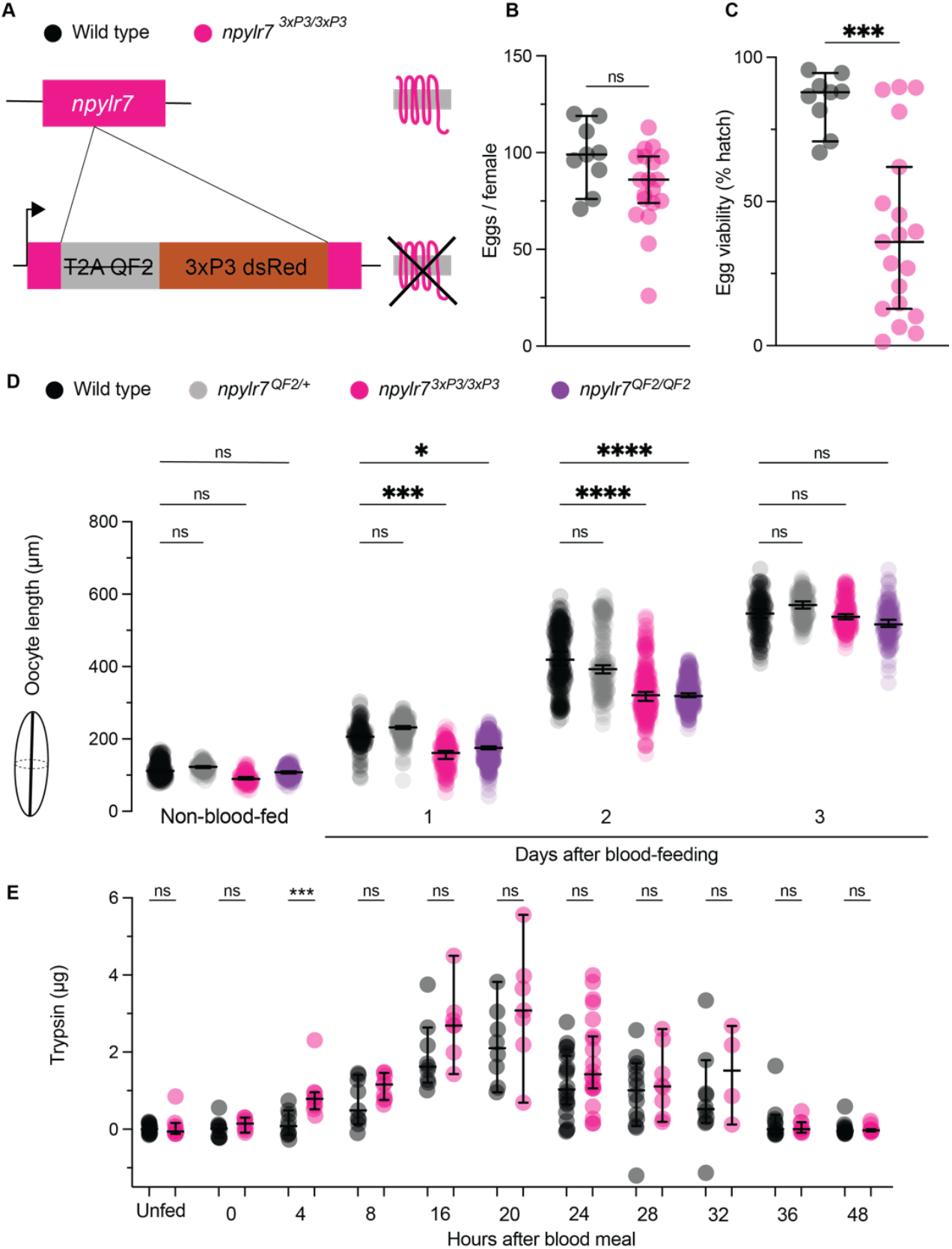
Oocyte provisioning and fecundity deficits in an independently-generated *npylr7* mutant line. (A)Schematic of *npylr7*^*3xp3*^ mutant allele. (B) Total eggs laid, n = 9-19 clutches from individual females (Unpaired *t*-test, ns = p > 0.05). (C) Egg viability (percent of eggs hatched per female), n = 9-19 clutches (from females of S2B) (Mann-Whitney test, *** = p < 0.001). (D) Oocyte length measured as distance from apical to basal end along longest axis using Fiji. n = 5-10 females with 10-20 oocyte measurements per ovary (Kruskal-Wallis test with Dunn”s multiple comparisons, * = p < 0.05, *** = p <0.001, **** = p < 0.0001). (E) Trypsin measurements in the gut. n = 4-10 female midguts (Mann-Whitney test, *** = p < 0.001, ns = p > 0.05). Data are shown as medians with 95% confidence intervals for visual consistency. Statistical comparisons were performed using parametric tests for normally distributed data and non-parametric tests for non-normal data, as determined by normality test (Shapiro-Wilk). Post-hoc multiple comparison tests were conducted against the wild-type control.

**Figure S3:**
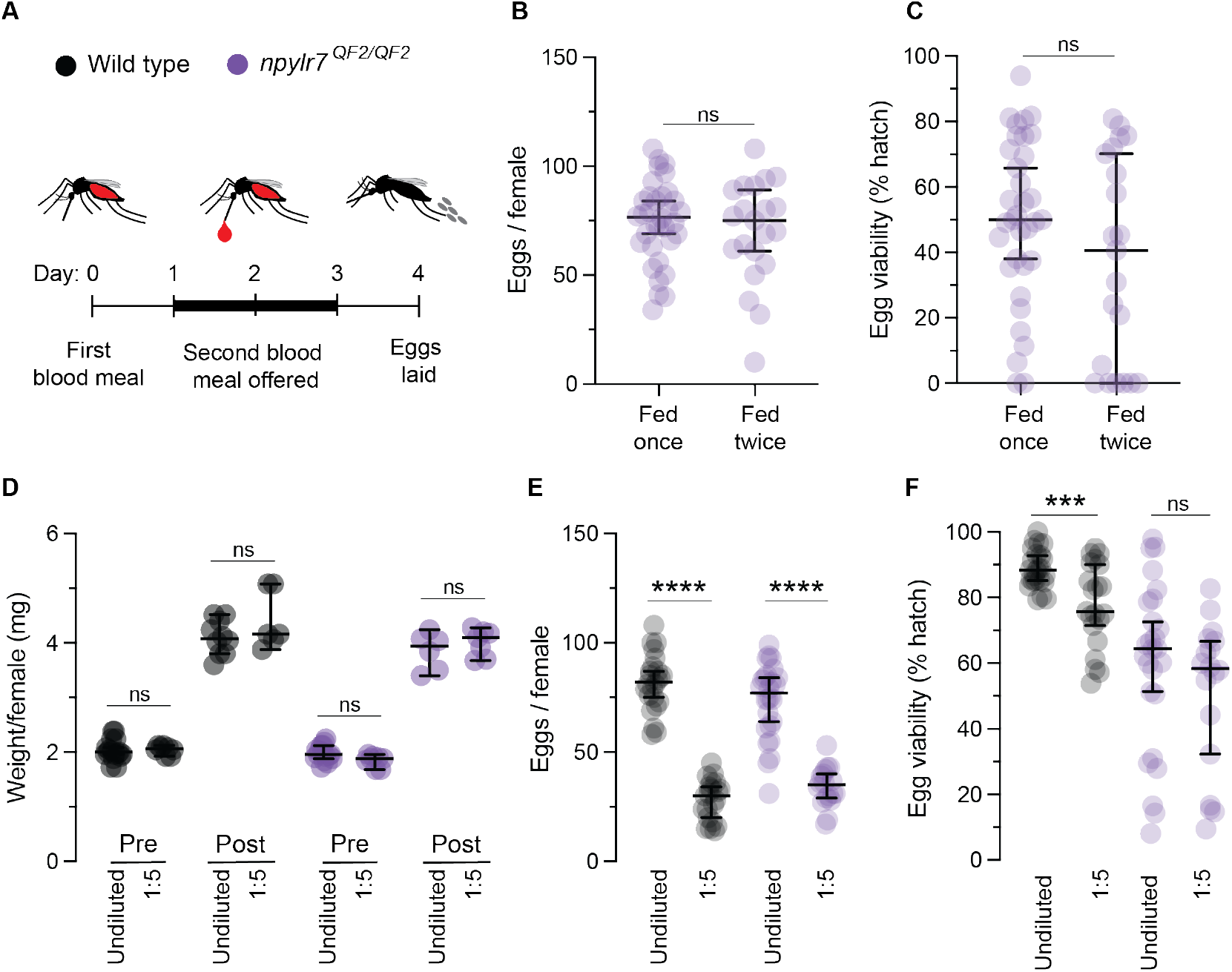
*npylr7* phenotypes are distinct from blood nutrient availability. (A) Timeline of refeeding assay in *npylr7* mutants. (B) Total eggs laid per female. n = 19-21 clutches from individual females (Unpaired *t*-test, ns = p > 0.05). (C) Egg viability (percent of eggs hatched per female) after refeeding. n = 19-21 clutches (from females of S3B) (Mann-Whitney test, ns = p > 0.05). (D) Weight per female after feeding on meals indicated (weighed in groups of 5, 7-17 groups) (Unpaired *t*-test for wild-type pre and *npylr7*^*QF2/QF2*^ post, Mann-Whitney test for wild-type post and *npylr7*^*QF2/QF2*^ pre. ns = p > 0.05). (E) Total eggs laid by individual females. n = 18-26 clutches (Unpaired *t*-test,**** = p < 0.0001). (F) Egg viability (percent of eggs hatched per female). n = 18-26 clutches. (Unpaired *t*-test for undiluted comparison, Mann-Whitney for 1:5 comparison, ns = p > 0.05, *** = p < 0.001). Data are shown as medians with 95% confidence intervals for visual consistency. Statistical comparisons were performed using parametric tests for normally distributed data and non-parametric tests for non-normal data, as determined by normality test (Shapiro-Wilk). Post-hoc multiple comparison tests were conducted against the wild-type control.

**Figure S4:**
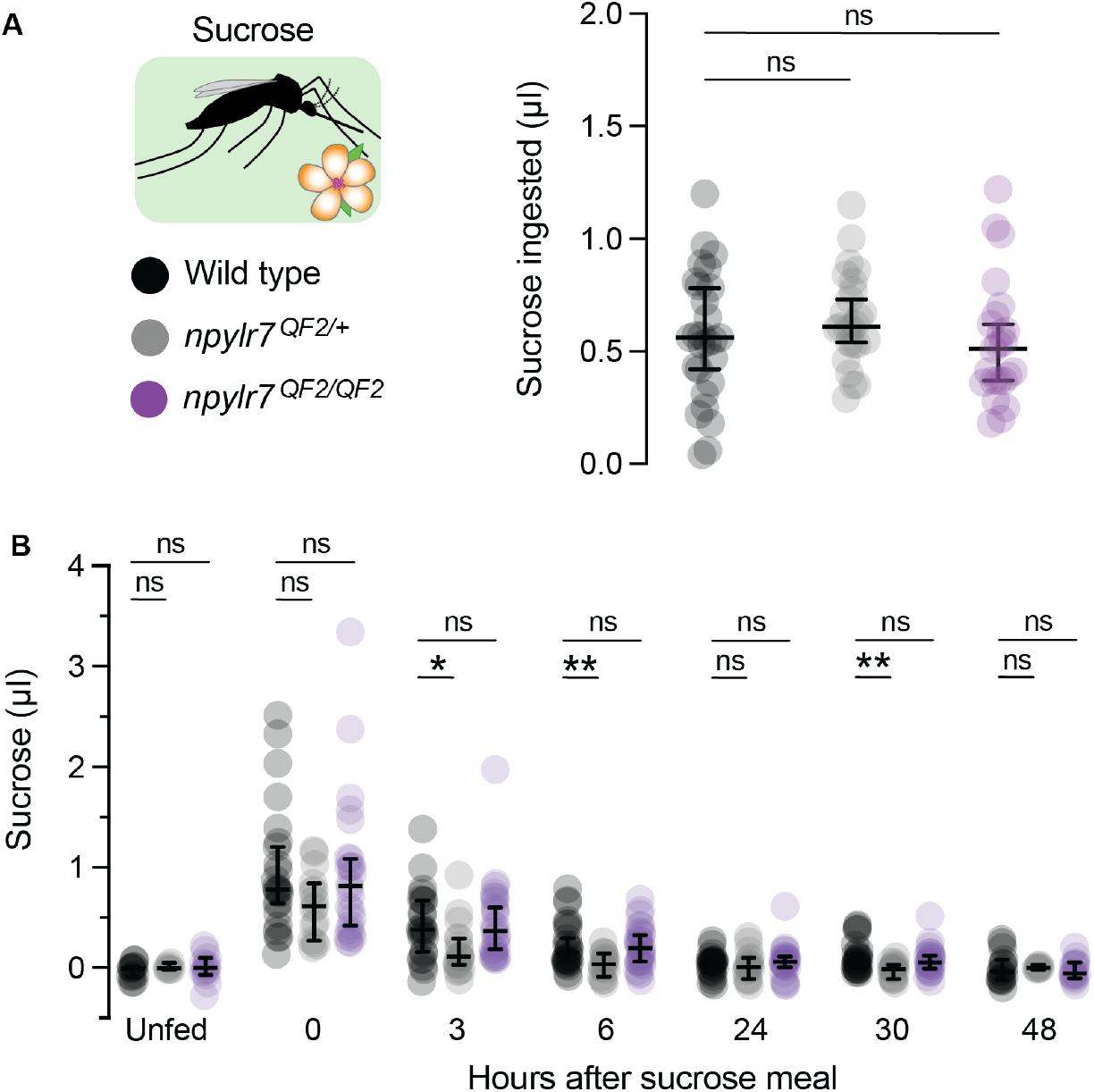
Sucrose consumption and clearance are unaffected in *npylr7* mutants. (A) Sucrose volume ingested, n = 23-27 females (Kruskal-Wallis test with Dunn”s multiple comparisons, ns = p > 0.05). (B) Sucrose clearance measured across time as amount of fluorescent sucrose meal retained, n = 6-27 females (Kruskal-Wallis test with Dunn”s multiple comparisons,ns = p> 0.05, * = p < 0.05, ** = p < 0.01). Data are shown as medians with 95% confidence intervals for visual consistency. Statistical comparisons were performed using parametric tests for normally distributed data and non-parametric tests for non-normal data, as determined by normality test (Shapiro-Wilk). Post-hoc multiple comparison tests were conducted against wild-type control.

**Figure S5:**
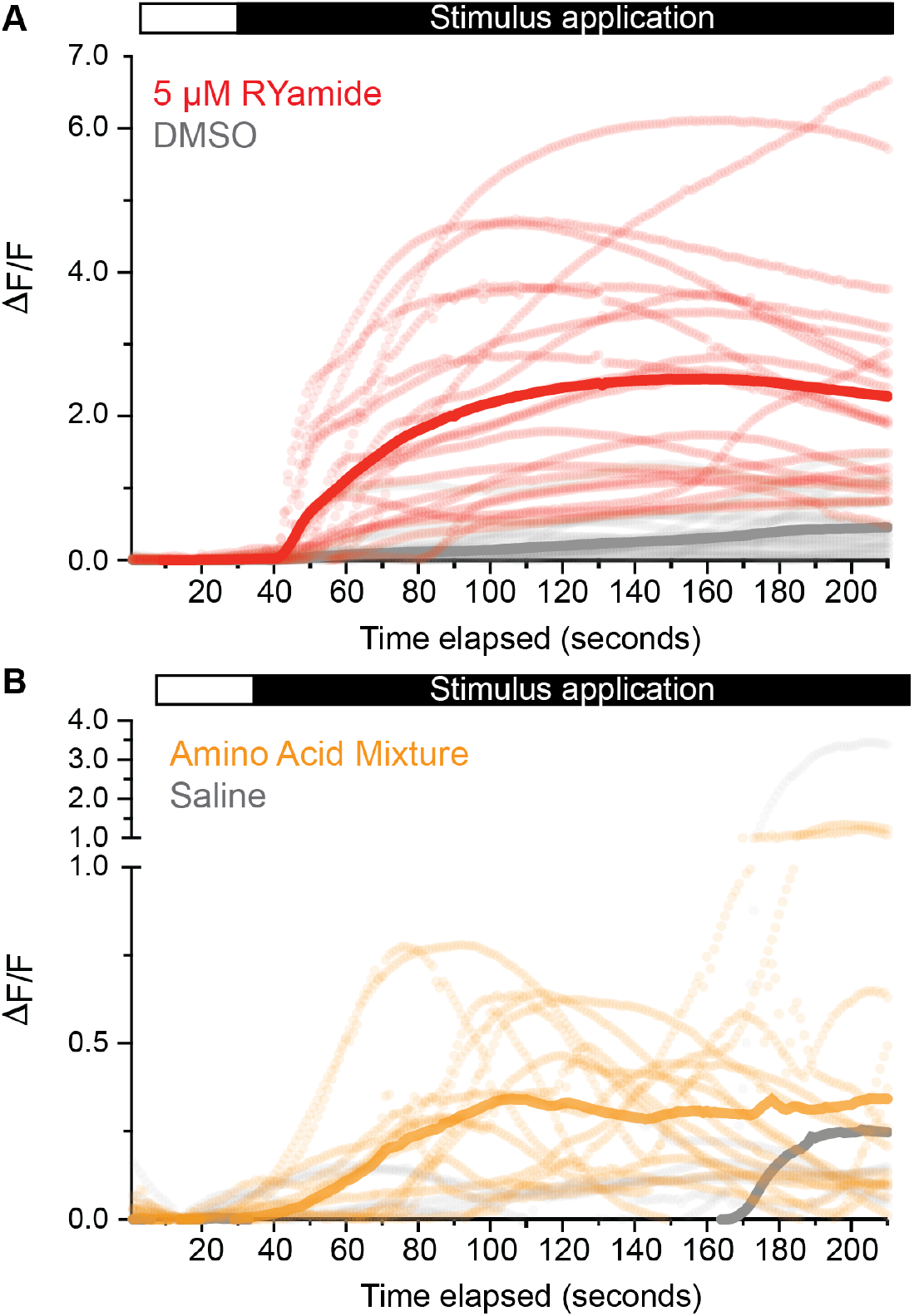
Raw traces of live imaging timecourse. (A) Live imaging timeline of ΔF/F after application of 5 *µM* RYamide peptide, or DMSO vehicle control. Mean is shown as dark line with individual replicates shown as dashed lines. Red = RYamide, gray = DMSO vehicle control. (B) Live imaging timeline of ΔF/F after application of amino acid mixture mimicking human blood or amino acid-free saline. Mean is shown as dark line with individual replicates. Yellow = amino acid mixture, gray = saline control.

